# Transcriptomic reconstruction of an interregional molecular connectome in schizophrenia

**DOI:** 10.64898/2026.10.05.756648

**Authors:** Ohhyeon Kwon, Uiyun Kim, Maree J. Webster, Sanghyeon Kim, Doheon Lee

## Abstract

**Background:** Schizophrenia is increasingly viewed as a disorder of multi-regional brain networks, but molecular and pathophysiological hypotheses are often studied at the level of single brain regions or neurotransmitter systems. A multi-regional, multi-neurotransmitter analysis is needed to clarify how molecular abnormalities relate to circuit-level hypotheses of the disorder.

**Methods:** Transcriptomic data from 2,209 post-mortem brain samples spanning five brain regions in 1,037 individuals were integrated. Directed connectivity strengths between regions were estimated using co-expression between presynaptic and postsynaptic neurotransmission-related genes. These estimates constitute an “interregional molecular connectome”. Macaque transcriptomic data were used to benchmark the estimation pipeline against reference macaque connectomes. Connectivity differences between healthy controls and patients with schizophrenia were calculated, and a directional consistency filter was applied to identify candidate dysregulations. An LLM-augmented literature screening framework was used to assess the level of literature support for our candidates.

**Results:** Our estimation pipeline showed concordance with reference connectomes, outperformed whole-transcriptome and random-gene co-expression measures, and remained stable when applied to macaque data. A total of 10 connections were marked as candidate dysregulations in schizophrenia at a nominal permutation threshold. Three of the candidates align with established anatomical connectivity, and another three with anatomical connections through relay regions. These candidates were characterized by bidirectional abnormalities in the cortico-striatal circuitry. The remaining four candidates had no well-established monosynaptic connection between two regions. These candidates were interpreted as co-regulated co-expression coordination changes. Literature screening quantified the number of articles discussing each candidate and how many of the articles directly support or refute each candidate. Our candidates collectively highlight the role of the cortico-striato-thalamo-cortical circuit in the pathophysiology of schizophrenia.

**Conclusions:** Interregional co-expression of neurotransmission-related genes carries more anatomical information than whole-transcriptome co-expression. Applied to schizophrenia, our analysis recapitulates known hypotheses and also proposes hypothesis-generating data-driven findings for experimental follow-up. This approach complements conventional region-wise analyses. Our approach could support a unified, multi-regional, and multi-neurotransmitter perspective of schizophrenia and point to candidate molecular mechanisms for further studies.

## Background

Schizophrenia is a severe psychiatric disorder characterized by profound and heterogeneous symptoms, including psychosis and cognitive impairment[1]. It is also highly heritable, with twin-study estimates of approximately 80%[2], and imposes a substantial burden on global public health[3]. Antipsychotics have predominantly targeted the dopamine D2 receptor (DRD2), providing partial alleviation of positive symptoms[4] while leaving negative and cognitive symptoms largely unaddressed[5]. An incomplete understanding of the molecular mechanisms underlying schizophrenia hinders the development of therapeutics with improved efficacy and fewer side effects[6]. Although large-scale genome-wide association studies have mapped hundreds of risk loci[7], translating these genetic discoveries into a mechanistic understanding of pathophysiology remains a major challenge[8]. As a consequence, an explanation that connects molecular findings to the clinical symptoms of the disorder is required.

Extensive research on molecular, pathological, morphological, and physiological endophenotypes has been directed toward closing this gap[9]. These efforts have led to diverse pathophysiological models, including two models that broadly organize current understanding of the disorder[10]. One is a neurochemical model built on molecular abnormalities[11]. The other is a neural disconnection model[12], which is built on emerging evidence of aberrant neurotransmission and network disconnectivity at both macrostructural and microstructural levels[13, 14]. Dopamine is strongly implicated in the neurochemical model and is one of the most widely studied neurotransmitters in the pathophysiology of schizophrenia[15]. Cumulative evidence implicates multiple neurotransmitters, including glutamate[7, 15], serotonin[16], γ-aminobutyric acid (GABA)[11], acetylcholine[17], and glycine[18]. Dysregulated neurotransmission and synaptic pathology are closely related, as suggested by the synaptic hypothesis of schizophrenia[19]. Multiple lines of evidence support these associations, including abnormal neurotransmitter levels[20], disrupted receptor function[21], and impaired neurotransmitter metabolism[22]. Neuroimaging evidence of functional disconnections among brain networks additionally illustrates the multi-regional and heterogeneous characteristics of the disorder[23].

Despite extensive study, results remain highly inconsistent, even within the same brain region. Methodological discrepancies between studies, uncontrolled confounding variables, and profound clinical heterogeneity across cohorts leave individual studies underpowered to resolve a consistent molecular signal[24]. In addition, prior investigations have largely focused on individual neuroanatomical regions. As a result, a disorder with pathology demonstrated across interacting brain regions has been characterized one region at a time. These two limitations constrain our understanding of schizophrenia as a systemic, multi-regional disorder. Each limitation requires a different strategy. The first is to combine cohorts at a scale large enough to account for confounding variables, rather than trying to avoid them. The second is to estimate quantities defined between regions, not within a single region. A single-region study cannot estimate such quantities, no matter how large the sample size becomes. Therefore, a “multi-regional, multi-neurotransmitter” approach is needed to determine how molecular disruptions relate to interregional connectivity changes. That approach could also identify candidate mechanisms for experimental follow-up.

Gene expression offers one route to that goal. Interregional gene co-expression covaries with anatomical and functional connectivity in the human and rodent brain[25]. This covariation supports the use of expression as a proxy for estimating connectivity beyond its role as a simple correlate. Analyses relating gene expression to brain connectivity have been applied to in vivo resting-state functional connectivity in the healthy brain[26, 27], and to structural dysconnectivity in schizophrenia[28]. A practical constraint shapes those designs. A human brain transcriptome requires tissue and is therefore obtained post mortem, whereas connectivity measurement in vivo requires a living subject. The two measurements cannot be acquired from the same brain at the scales required for their respective analyses. Studies relating them have paired an expression atlas from one set of donors with connectivity from different cohorts, and correlated the two across anatomical space[26–28]. In this pairing, expression is treated as a fixed anatomical reference, and individual variation is averaged into the atlas. Diagnostic differences are represented only in the connectivity data[28].

An alternative approach is to derive both sides of the correlation from expression measured across regions within the same individuals. In this approach, diagnosis becomes a property of the samples themselves, and an anatomical reference enters only as an external benchmark. Statistical methods to infer connectivity from expression alone already exist. Several approaches infer cell-cell communication strength from ligand-receptor expression profiles using single-cell RNA sequencing (scRNA-seq) data[29, 30]. A similar approach has also been proposed to infer communication between neural regions based on neurotransmission-related genes[31]. However, these approaches have not been applied to a case-control post-mortem cohort, because they were designed for single-cell data. The need for single-cell data constrains the cohort size and the regional coverage available for such a comparison. To our knowledge, a neurotransmitter-resolved reconstruction of interregional coordination between presynaptic and postsynaptic gene expression using such a cohort has not been reported.

An analysis that spans multiple region pairs and neurotransmitter systems generates more than a hundred testable hypotheses. Systematically evaluating high-dimensional predictions against the vast unstructured biomedical literature remains difficult. The volume of research is expanding rapidly, with PubMed alone accumulating about 1.7 million new articles in 2025[32], making exhaustive manual review impractical. Recent advances in large language models (LLMs), particularly their reasoning capabilities and natural language comprehension[33], offer a practical solution for screening large volumes of biomedical literature[34]. Vector-based similarity search with retrieval-augmented generation (RAG) is now common. This approach relies on the semantic similarity between compressed representations for information aggregation[35]. Similarity-based search prioritizes dominant patterns, while under-representing minor, conflicting signals, thus obscuring variance and heterogeneity across the literature. As a result, the method might over-generalize, under-report conflicting evidence, and produce unsupported claims when evaluating complex scientific hypotheses[36]. We addressed this through an LLM-augmented framework, adapted from our prior work[37]. The framework systematically parses and evaluates unstructured biomedical literature by assessing each article separately. The framework classifies individual articles as supporting, refuting, or neutral with respect to each predicted dysregulation. Article-level classification allows findings to be systematically stratified by the degree of literature evidence based on the proportion and volume of retrieved articles classified as supporting or refuting.

Here, we reconstruct a directed neurotransmission connectivity network, which we call an “interregional molecular connectome”, from post-mortem transcriptomic data to investigate the multi-regional molecular pathophysiology of schizophrenia (**Fig. 1A**). We use “connectome” here for a network of coordinated transcription between regions, and the direction of an edge follows from the presynaptic or postsynaptic role of the participating genes, not from a measured projection. Connectivity strength between two regions is estimated from interregional co-expression between presynaptic and postsynaptic neurotransmission-related genes. The co-expression measure is background-corrected to mitigate the effect of inherent correlation, and weighted by the estimated abundance of the corresponding neurotransmitter of the source region (**Fig. 1B**). We used macaque transcriptomic data to benchmark the connectivity estimation performance against two reference connectomes from tract-tracing analysis. We integrated seven transcriptomic datasets from patients with schizophrenia and healthy controls, comprising 2,209 brain samples from five brain regions across 1,037 individuals. Connectivity changes in schizophrenia were identified by comparing estimated strengths between diagnostic groups. Label permutation analysis followed by a directional consistency filter was applied to the changes to identify candidate dysregulations. These candidates were interpreted by prioritizing genes that drove the changes and analyzing specific molecular mechanisms related to the prioritized genes. We stratified candidates by established interregional connections. Most candidates fell on well-established connections or indirect connections through relay regions. These candidates aligned with the cortico-striato-thalamo-cortical (CSTC) circuit. The remainder did not align with established interregional connections, which may reflect shared afferent innervation or a common regulatory axis. To synthesize evidence for our findings, we applied an LLM-augmented framework for literature screening. The screening evaluated our results against the relevant biomedical corpus to quantify the degree of literature support for our data-driven hypotheses. By estimating neurotransmitter-specific coordination between regions within the same individuals, this work connects the neurochemical and disconnection models that have developed largely in parallel. Consequently, this work suggests a multi-level view of the neurotransmission dysfunction underlying schizophrenia.

**Fig. 1.**
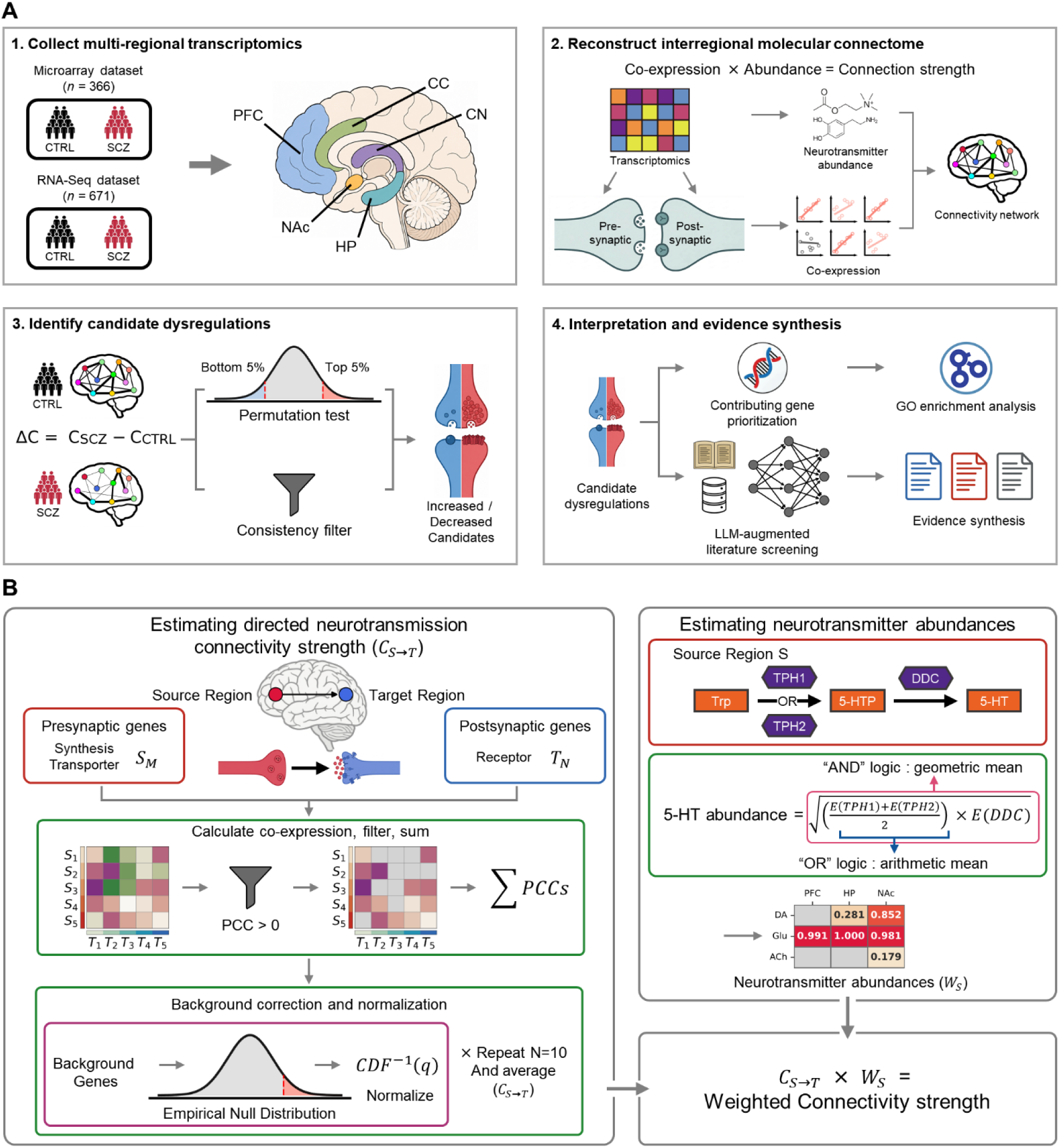
Study design and reconstruction of the “interregional molecular connectome”. (**A**) Overall workflow. (**B**) Estimation of directed neurotransmission connectivity from a source region S to target region T for one neurotransmitter system, to reconstruct an “interregional molecular connectome”.

## Methods

### Human transcriptomic data collection and preprocessing

A total of seven transcriptomic datasets, comprising two microarray (MA) and five RNA sequencing (RNA-seq) datasets[38–43], were integrated to support cross-platform analysis. Datasets were curated from public repositories, with selection based on the availability of samples from individuals with schizophrenia and from control individuals. Datasets included samples from five brain regions: prefrontal cortex (PFC), cingulate cortex (CC), hippocampus (HP), caudate nucleus (CN), and nucleus accumbens (NAc). We manually inspected demographic variables for each dataset and verified genomic sample identity using ‘NGSCheckMate’[44] for RNA-seq datasets, to identify and exclude potential duplicate samples across studies. Samples from individuals under 18 years of age were then excluded, following guidelines from the ‘BrainSeq’ Consortium[40]. After quality control, the final cohort comprised 201 controls and 165 patients with schizophrenia from MA datasets, and 389 controls and 282 patients with schizophrenia from RNA-seq datasets. A total of 2,209 brain samples were obtained from 1,037 donors. Interregional co-expression could be estimated only for donors with samples from at least two brain regions. Thus, 1,800 samples from 628 donors were included in the interregional analysis, and the number of available donors varies across region pairs. The remaining 409 donors each contributed a single sample, which was used only for abundance estimation. Detailed demographic data, accession numbers, and region availability for each dataset are provided in **Table 1 and Supplementary Table S1**. All datasets were obtained with approval from the relevant ethics committees for the original projects.

**Table 1.** Demographic characteristics of the seven human post-mortem transcriptomic datasets.

| Dataset name | Assay type | Sampled regions | Diagnosis | Sex | Age (range, mean) | Accession | Reference |
| --- | --- | --- | --- | --- | --- | --- | --- |
| GSE53987 | MA | PFC, HP, CN | 19 CTRL, 19 SCZ | 20 M, 18 F | 22.0-68.0 (46.6) | GEO: GSE53987 | [38] |
| HBCC | MA | PFC, HP | 182 CTRL, 146 SCZ | 218 M, 110 F | 20.2-97.0 (49.0) | dbGaP: phs000979.v3.p2 | [43] |
| Array Collection | RNA-Seq | PFC, HP, CC, CN, NAc | 35 CTRL, 35 SCZ | 52 M, 18 F | 19.0-60.0 (43.4) | Controlled Access; inquire SMRI | [42] |
| New Collection | RNA-Seq | PFC, HP | 19 CTRL, 20 SCZ | 29 M, 10 F | 28.0-70.0 (49.3) | Controlled Access; inquire SMRI | [42] |
| Neuropathology Consortium | RNA-Seq | PFC, HP, CC | 15 CTRL, 14 SCZ | 18 M, 11 F | 25.0-68.0 (45.9) | Controlled Access; inquire SMRI | [42] |
| GSE80655 | RNA-Seq | PFC, CC, NAc | 24 CTRL, 24 SCZ | 42 M, 6 F | 24.0-70.0 (46.5) | GEO: GSE80655 | [39] |
| BrainSeq | RNA-Seq | PFC, HP, CN | 296 CTRL, 189 SCZ | 338 M, 147 F | 18.0-96.9 (48.3) | PFC, HP: public, via BrainSeq homepage; CN: dbGaP: phs003495.v1.p1.c1 | [40, 41] |
| Total | MA, RNA-Seq | PFC, HP, CC, CN, NAc | 590 CTRL, 447 SCZ | 717 M, 320 F | - | Public, Controlled |  |

Counts are donors after removal of duplicate samples and exclusion of donors under 18 years of age.
Age is given in years.

A unified preprocessing pipeline was applied to integrate expression profiles across platforms. For microarray data, raw intensity files were processed using the ‘affy’[45] and ‘beadarray’[46] R packages for Affymetrix and Illumina platforms, respectively. Probe-level intensities were then normalized and collapsed to gene-level expression values using annotations from the ‘biomaRt’[47] and ‘illuminaHumanv4.db’[48] packages. For RNA-seq data, raw sequencing reads were first checked for their quality using ‘FastQC’[49], followed by adapter trimming and low-quality read filtering using ‘Trimmomatic’[50]. Next, reads were quantified against the human genome reference ‘GRCh38.p14’[51] using the pseudo-alignment tool ‘Salmon’[52]. Transcript-level quantifications were aggregated to gene-level counts using the ‘tximport’[53] R package. An expression threshold following the GTEx consortium standards[54] (retaining genes with > 0.1 transcripts per million and ≥ 6 reads in at least 20% of the samples) was applied to filter out noise from low-expression genes. The count data were converted to counts per million (CPM) values and log2-transformed by adding 0.25 (log2(CPM + 0.25)), following the approach of Song *et al*.[55]. Finally, we applied surrogate variable analysis (SVA) using the ‘sva’[56] R package to mitigate batch effects and other technical variations. The SVA correction removed the effect of latent variables, retaining the disorder- and region-specific variation at the same time (**Supplementary Fig. S1**).

### Curation and functional classification of neurotransmission genes

A comprehensive list of genes involved in synaptic transmission was manually curated to estimate neurotransmission connectivity strength between regions. The gene list was initially aggregated from four established sources: 1) the Allen Human Brain Atlas[57] neurotransmitter-related gene set (in the ‘Gene list’ file)[58]; 2) the comprehensive macaque transcriptomic resource by Bo *et al*.[59]; 3) the ‘Reactome’ database[60] (V94, accessed 2025-09-16) with selected pathway terms related to neurotransmission; and 4) the “NeuronChatDB”[31]. The initial aggregation yielded a superset of 585 genes (**Supplementary Information S1**). Each gene was manually inspected to ensure biological relevance and classified into one of eight distinct neurotransmitter systems: dopamine, serotonin, glutamate, GABA, acetylcholine, norepinephrine (noradrenaline), epinephrine (adrenaline), and glycine. Genes related to synaptic adhesion (specifically, neurexins and neuroligins) were included under a separate synaptic adhesion category. For each system, genes were subdivided into two functional categories, a distinction required for modeling directed connectivity:

- “Category I” (signal transduction, postsynaptic): Genes encoding receptors, ion channels, and downstream effectors that are required for postsynaptic signal reception.
- “Category II” (metabolism and transport, presynaptic): Genes encoding synthesis and catabolism enzymes, vesicular transporters, and reuptake transporters that are required for presynaptic signal generation and termination.

After the curation, 151 genes related to eight neurotransmitters and synaptic adhesion were retained for analysis as the “curated gene list” (**Supplementary Table S2**). Genes falling below the expression threshold in a given region were excluded from that region’s analysis.

### Reconstruction of neurotransmission connectivity networks

We developed an approach that estimates the directed connectivity strength from a source region S to a target region T using expression levels of genes related to neurotransmission (**Fig. 1B**). We assumed that directed signaling strength is reflected by correlations between presynaptic Category II gene expression in source region S and postsynaptic Category I gene expression in target region T, for each neurotransmitter system. For a given neurotransmitter system with M Category II genes {*S*_1_, *S*_2_, …, *S_M_*} in the source region and N Category I genes {*T*_1_, *T*_2_, …, *T_N_*} in the target region, we calculated the Pearson correlation coefficients (PCCs) for every possible interregional gene pair {(*S*_1_, *T*_1_), (*S*_1_, *T*_2_), (*S*_1_, *T*_3_), ⋯, (*S_M_*, *T_N_*)}. The pairwise calculation yielded an M-by-N correlation coefficient matrix for each region pair. Under our assumption that positive co-expression reflects coordinated regulation, we summed only the positive PCCs. The “summed PCCs” served as an uncorrected estimate of the connectivity strength of neurotransmission between S and T.

To define the connectivity strength and control for inherent correlation, we applied a background correction step. For a neurotransmitter system with M Category II genes and N Category I genes, we randomly sampled M+N genes from the background gene panel and calculated the summation of positive PCCs. The background gene panel consisted of 585 randomly drawn genes from the candidate background gene pool, matching the size of the initial neurotransmission-related superset. The candidate background gene pool consisted of genes that were expressed in all regions, excluding the 585 initially aggregated neurotransmission-related genes. Gene sampling was repeated 1,000 times, and the summed PCCs of sampled genes formed an empirical null distribution. The observed sum of PCCs for a given neurotransmitter system was then converted to a quantile value relative to this null distribution. This quantile was transformed using the inverse cumulative distribution function (*CDF*^−1^) of the standard normal distribution to generate a normalized z-score. The background correction was repeated across 10 background gene panels (together, a ‘background gene panel set’) to reduce dependence on any single panel. The average of the 10 z-scores constituted the final connectivity strength from the source region S to the target region T (*C_S_*_→*T*_). Throughout, we write a connection from source region S to target region T as S → T.

### Weighting connectivity strengths by neurotransmitter abundances

Although gene expression correlation reflects potential signaling strength, physiological signal transmission strength also depends on the availability of the neurotransmitter, because neurotransmitter levels and neurotransmission capacity are correlated[61]. To incorporate this, we estimated the regional neurotransmitter abundance for each source region using the “AND-OR” logic adapted from the “NeuronChat”[31] method and multiplied each abundance estimate by the corresponding connectivity strength (*C_S_*_→*T*_). Briefly, this logic models the biosynthetic pathway of each neurotransmitter:

- “AND” logic: For a complete metabolic pathway synthesizing a neurotransmitter, estimating neurotransmitter abundance requires the joint expression of all participating enzymatic genes in the pathway.
- “OR” logic: For enzymes catalyzing the same metabolic step, any one of them can handle it, allowing the metabolic step to proceed through alternative routes.

The AND logic was applied using the geometric mean of expression levels, and the OR logic was applied using the arithmetic mean. Both rules were applied as described in the original “NeuronChat” study, with more details available in the original publication[31]. Gene functions for Category II genes were annotated using the ‘GeneCards’ database[62], and the annotations were used to define the core metabolic pathway for each neurotransmitter. Two enzymes absent from our curated gene list (PAH and GLUD1) were added to the metabolic pathway because they participate in key metabolic processes. For each neurotransmitter, we estimated its abundance (*W_S_*) using expression levels of enzymes from each region based on the pathway, applying the “AND-OR” logic. We assumed that the amount of the initial metabolic substrate was not limiting. For synaptic adhesion, the abundance was defined as the mean expression level of neurexin genes (NRXN1, NRXN2, and NRXN3). Abundance estimates were calculated for each sample, and group means were used in weighting connectivity strengths, as our estimator is defined on group-level correlations. The final weighted connectivity strength (*C^W^*) was defined as the product of the averaged connectivity strength (*C_S_*_→*T*_) and the group mean of estimated neurotransmitter abundance (*W_S_*) in the source region S. This weighting of connectivity strengths incorporates the estimated synthesis capacity of the source region into the co-expression-based measure.

### Benchmarking against macaque tract-tracing-derived connectomes

To evaluate the biological accuracy of our network reconstruction, we benchmarked our method against reference anatomical connectomes from macaque tract-tracing data. Two macaque connectome datasets derived from the tract-tracing database “CoCoMac”[63–65] were used:

- ‘Organized CoCoMac’: A manually curated set of directed connections derived from tract-tracing analyses, organized hierarchically by anatomical subdivision using the “CoCoMac” database[66].
- ‘TheVirtualBrain’[67]: A connectome incorporating neuroimaging-derived weights and refined parcellations[68], also using the “CoCoMac” tract-tracing database.

We explicitly note that the two reference connectomes are not independent. The connectomes derive from the CoCoMac tract-tracing database using different anatomical mappings, and TheVirtualBrain connectome carries additional imaging-derived edge weights that cannot resolve the direction of a connection. Transcriptomic data for benchmarking were obtained from GSE180092, which provides high-resolution bulk RNA-seq data from 36 macaques, covering 15 cortical and subcortical regions per animal[69]. These transcriptomic regions were manually mapped onto the anatomical parcellations used in both connectome datasets (**Supplementary Table S3**). The Organized CoCoMac connectome does not distinguish brain hemispheres, while TheVirtualBrain connectome has separate left and right hemispheres. Since the transcriptomic data were only taken from the right hemisphere, we used the right-hemisphere portion of TheVirtualBrain connectome. Self-loops originating from subregion merging were not used for benchmarking analysis. Our connectivity reconstruction method was applied to the macaque transcriptomic data to infer interregional neurotransmitter-specific connectivity strengths. For each region pair, we summed the eight neurotransmitter-specific connectivity strengths and synaptic adhesion strength to obtain a single score. The predicted strength was compared against the two reference connectome datasets. Performance was quantified as the area under the receiver operating characteristic curve (auROC) and the area under the precision-recall curve (auPRC). Each neurotransmitter-specific connectivity estimate was used to calculate auROC and auPRC values against the two connectomes. Because monoamine neurotransmitters (dopamine, serotonin, norepinephrine, and epinephrine) have zero estimated abundance in five of the macaque cortical regions (**Supplementary Fig. S2**), we also recalculated the neurotransmitter-specific benchmarking metrics with the predicted connections and both reference connectomes restricted to the 10 remaining regions. Acetylcholine was excluded from neurotransmitter-specific benchmarking because the metrics were not derivable for either connectome. In the Organized CoCoMac connectome, three regions with nonzero estimated acetylcholine abundance have only a single positive reference edge. In TheVirtualBrain connectome, no reference edges are defined for these regions.

The predictive performance measures of our method were compared with those from the “NeuronChat” method. To test whether the “NeuronChat” algorithm can be generalized to bulk RNA-seq data, we included the pseudo-bulk-based reconstruction of the experimentally validated mouse brain connectome. For benchmarking, we used the same datasets as the original “NeuronChat” study (retrograde injection analysis data from Tasic *et al*.[70] and scRNA-seq data from GSE115746[70] and GSE185862[71]). The ‘Seurat’[72] R package was applied to aggregate the single-cell-level transcriptomic data into pseudo-bulk expression profiles using the mouse sample information provided in the original study. After the aggregation, we applied our preprocessing pipeline to generate processed sample-by-gene expression matrices and ran the original “NeuronChat” package to infer the regional connectome. We then calculated auROC and auPRC for the primary visual cortex (VISp) and anterior lateral motor cortex (ALM) models using the inferred connection strength for each model against the experimentally validated retrograde injection analysis data. The “NeuronChat” package was used to infer connection strength using macaque transcriptomic data, and auROC and auPRC were calculated against the two reference macaque connectome datasets.

Auxiliary measures of connectivity estimates were additionally analyzed to assess the robustness of our method. First, we applied bootstrap resampling 1,000 times and calculated auROC and auPRC for each of the resamples. Using the 1,000 resamples and their predictive performance measures, we calculated 95% confidence intervals (CIs) for each point estimate. Second, to identify whether the predictive power of the connectivity estimates came from the inherent interregional co-expression or from co-expression coordination between neurotransmission genes, we designed gene ablation analyses. The sum of interregional PCCs for every gene pair served as a connectivity strength estimate between regions. We also repeated our connectivity strength estimation for 1,000 randomly selected gene sets that matched the curated gene list in size and expression characteristics, i.e., expression decile and whether the gene was expressed in the region. The connectivity estimates from these two ablations were used to calculate auROC and auPRC for the two reference connectome datasets. Third, we subsampled the macaque cohort 100 times at each sample size (n = 8, 12, …, 32), sampling without replacement. We then calculated connectivity estimates and derived auROC and auPRC for each of the resamples. Fourth, each of the 100 subsamples was randomly split into two halves. At n = 36, the entire cohort was randomly split into two halves 100 times, without subsampling. For each half, connectivity strengths were estimated, and Spearman’s rank correlation (ρ) of connectivity estimates between the two halves was calculated. Finally, we repeated the full pipeline across different background gene draws and quantified predictive performance against the macaque connectomes to test whether connectivity reconstruction was robust to the choice of candidate background gene pool. The candidate background gene pools were constructed under two eligibility rules: 1) all genes expressed in both macaque and human expression profiles, and 2) the same pool after removing the 585 neurotransmission-related genes. For each eligibility rule, we drew four random background gene panel sets (10 panels × 585 genes = 5,850 genes in each panel set). Consequently, eight background gene panel sets were drawn, and we repeated our analysis using all eight.

### Identification of candidate dysregulations and biological interpretation

To identify connectivity dysregulation in schizophrenia, we compared the weighted connectivity strengths between the patients with schizophrenia and control groups. The difference score (Δ*C^W^*) was defined as the weighted strength estimate of controls subtracted from that of patients with schizophrenia 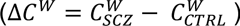. A positive difference indicates increased connectivity, and a negative difference indicates decreased connectivity in patients with schizophrenia. Statistical significance was assessed by a non-parametric permutation test. The diagnostic labels (schizophrenia/control) were randomly shuffled 10,000 times. Δ*C^W^* was calculated by recomputing *C_S_*_→*T*_ and *W_S_* for each permutation to construct an empirical null distribution of difference scores Δ*C^W^*. A connection was considered nominally significant if its observed difference fell within the one-tailed 5% tail (upper or lower 5%) of the empirical null distribution (label-permutation quantile < 0.05 or > 0.95, which is equivalent to a two-tailed 0.1). For each connection, we computed two-tailed nominal permutation p-values using its empirical null distribution, together with Benjamini-Hochberg (BH)-adjusted q-values across all tested connections. Candidates were selected using the nominal threshold, with corresponding p-values and q-values reported in **Supplementary Table S4** for reference.

Because the weighted connectivity reflects both altered co-expression coordination and altered neurotransmitter abundance, we compared abundance between diagnostic groups in each source region. A two-tailed t-test was used to quantify the statistical significance of group differences. Group difference effect size was quantified by Hedges’ g. A directional consistency filter was then applied. An increase in connectivity (Δ*C^W^* > 0) was retained if the estimated neurotransmitter abundance in the source region was increased in patients with schizophrenia, or at least not significantly decreased (t-test p > 0.05). Conversely, a decrease in connectivity (Δ*C^W^* < 0) was retained if the estimated neurotransmitter abundance in the source region was decreased, or at least not significantly increased (t-test p > 0.05). This filter ensured that the direction of the connectivity strength difference did not contradict the direction of estimated abundance difference in the source region. This step is an internal consistency check, not independent verification. Connections that fell in either of the 5% tails (top or bottom) and passed the directional consistency filter were retained as candidate dysregulations. Additionally, the standardized connectivity difference ((Δ*C^W^* - null distribution mean) / null distribution standard deviation) was calculated to indicate the relative strength of difference. The standardized difference is reported as a descriptor of relative magnitude only. Permutation quantiles were the only numeric values used in deriving nominal significance, and the standardized differences were not converted to p-values. The effect size of group difference (Hedges’ g) and the standardized difference help distinguish whether a dysregulation is driven by altered co-expression coordination or by changes in neurotransmitter abundance estimates.

As in the macaque benchmarking analysis, we additionally included auxiliary measures for the human cohort. First, we applied bootstrap resampling 1,000 times, calculated the difference score between two diagnostic groups for each resample, and calculated 95% CIs for our point-estimate difference scores along with the ‘sign stability’ that indicates the proportion of resampled differences that retained the sign of the point estimate. Second, we split the cohort in half 10 times, preserving dataset structure and diagnosis label composition. For each half, diagnostic group-level connectivity estimates and their difference scores were calculated, and Spearman’s ρ of the estimates and differences between the two halves was calculated for each of the 10 splits. For each of the 20 halves, the diagnostic labels were shuffled 500 times. For each label shuffle, we calculated the difference scores and then computed Spearman’s ρ from those differences. The shuffled difference scores gave an empirical null distribution with permutation p-value for each connection in that half. The shuffled Spearman’s ρ of difference scores between halves gave an empirical null distribution for half-split agreement. The smallest region pair comprised 25 controls and 29 patients with schizophrenia (**Supplementary Table S1**), yielding a minimum of 12 donors per diagnostic group in each half, so that all connections could be calculated in any split. Third, we removed each contributing dataset in turn and recalculated the difference scores. Region pairs had one to six contributing datasets (**Supplementary Table S1**), giving two to six leave-one-out estimates for pairs with at least two datasets. Region pairs with a single dataset could not be evaluated. Finally, we repeated the diagnostic label shuffle analysis under both candidate background gene eligibility rules (see “Benchmarking against macaque tract-tracing-derived connectomes” section for more details). To assess whether the identified dysregulations depended on the choice of eligibility rules, we compared the results from the two analyses as a sensitivity analysis.

For the candidate dysregulated connections, we prioritized genes that drive molecular changes. We first calculated the “correlation contribution” of each gene by summing the differences in PCC values within the dysregulated connections. Genes were ranked by their contribution, and the top 30% of genes were selected. The selected subset accounted for about 80% of the correlation contributions, as their distribution was concentrated near zero (**Supplementary Fig. S3**). The prioritized genes were then subjected to the Gene Ontology (GO) enrichment analysis (on the “Molecular Function” (GO:0003674), “Biological Process” (GO:0008150), and “Cellular Component” (GO:0005575) terms) using the PantherDB GO API (version 19.0)[73]. We first conducted the enrichment analysis using all the genes in the curated gene list, for each presynaptic/postsynaptic category and for each neurotransmitter system, and used these as the baseline. We next conducted the same analysis using only prioritized genes, and compared prioritized gene-specific enrichment results with baseline enrichment results by fold enrichment values and BH-adjusted q-values. Terms with elevated fold enrichment relative to baseline were selected to identify the specific molecular mechanisms underlying the connectivity dysregulations.

### Benchmarking LLM performance in literature screening

We applied an LLM-augmented literature screening framework that analyzed each article separately to synthesize evidence for our candidates, as similarity-based aggregation such as RAG can obscure minor or conflicting evidence. For this task, LLMs must be able to comprehend literature abstracts and determine whether a “premise” supports, refutes, or is neutral towards a given “hypothesis”, which is a task known as natural language inference (NLI)[74]. To establish the credibility of LLMs as a literature screening tool, we benchmarked their NLI performance on a biomedical-specific NLI benchmark dataset. We evaluated the selected model on the BioNLI dataset, which is a domain-specific curated benchmark with adversarial false examples that are designed to test NLI capability in biological hypothesis evaluation[75]. We used modified BioNLI data from our previous work, the ‘Believe’ framework[37]. The dataset contains 400 instances: 134 positive, 133 negative, and 133 neutral. Positive (true) data consist of paired abstracts and hypotheses where each hypothesis is a direct conclusion of its paired abstract. Negative data consist of paired abstracts and hypotheses, in which the hypothesis is lexically opposed to the abstract’s conclusion. Neutral data consist of pairs of unrelated hypotheses and abstracts, so no inference is licensed. Details of data modification are described in the original work[37].

For high-throughput literature screening, we selected “gpt-oss-120b”[76], which is an open-weight model developed by OpenAI. When using the model, we set ‘temperature’ = 0.0 and reasoning effort to ‘high’. Inference was run on in-house graphics processing units (GPUs). Multiple instances of “gpt-oss-120b” were executed on an in-house GPU server equipped with up to five NVIDIA RTX PRO 6000 Blackwell Server Edition GPUs, or eight NVIDIA A100 80GB GPUs. To benchmark our model choice, we included the best- and weakest-performing models evaluated in the ‘Believe’ framework[37], “gemini-3-pro-preview-11-2025-high” and “deepseek-v3.2-thinking”, for comparison with “gpt-oss-120b”. Both comparison models were accessed through the “OpenRouter” platform[77], with the ‘include_reasoning’ parameter set to ‘True’ and ‘temperature’ = 0.0 for the models. Each model was instructed to determine whether a given “abstract” supports, refutes, or is neutral towards a given “hypothesis”. The models were prompted to produce an output in a structured JSON format. After retrieving LLM outputs, we calculated accuracy, macro precision, macro recall, and macro F1 score[78] using the reference labels. Details of the system prompt and user prompt are provided in **Supplementary Information S2**.

### LLM-augmented literature screening framework

We first analyzed the feasibility of the literature screening framework using well-established biomedical hypotheses against literature abstracts. We manually wrote a set of such hypotheses and generated up to two adversarial negations for each hypothesis. Literature abstracts used in the screening process were retrieved from the PubMed[32] database (accessed 2026-08-10). PubTator 3.0[79] was used to normalize entity names and construct precise search queries for each given hypothesis. For each hypothesis, the abstracts returned by its query were fed into the literature screening framework to classify whether each abstract supports, refutes, or is neutral to each hypothesis. From the number of abstracts assigned to each verdict, we calculated the two literature screening metrics defined in the ‘Believe’ framework[37].

- Relevancy score = ((# Supports + # Refutes) / # Total abstracts), indicating the proportion of articles that discuss the hypothesis among retrieved articles. The number of total abstracts is the number of search results for the given query.
- Alignment score = (# Supports / (# Supports + # Refutes)), indicating the proportion of relevant retrieved articles classified as supportive.

After the feasibility analysis, we applied the literature screening framework to our candidate dysregulations. Our observations were converted into natural language hypotheses (e.g., “Glutamatergic signaling from the <SOURCE REGION> to the <TARGET REGION> is hyperfunctional in patients with schizophrenia”). Hypotheses were phrased in functional terms to match how these relationships are described in the literature, which improves retrieval. The phrasing does not imply that our estimates measure function. A uniform PubTator search query (@Disease_Schizophrenia) was applied when retrieving PubMed abstracts, and the retrieved abstracts were applied to all the hypotheses (total 246,074 abstracts, retrieved 2026-08-10). As a consequence, the total number of abstracts was identical across all hypotheses, and the Relevancy score was directly proportional to the number of articles classified as supporting or refuting. Candidate dysregulations were classified into three categories: 1) literature-supported (Alignment ≥ 0.5 and Relevancy > 0.0001, which is more than 24 articles), 2) controversial (Alignment < 0.5 and Relevancy > 0.0001), and 3) underexplored (Relevancy < 0.0001). Alignment scores were treated as unstable when 24 or fewer relevant articles were retrieved. These classification thresholds were set from the result of the feasibility analysis.

## Results

### Transcriptomic reconstruction shows concordance with reference anatomical connectivity

To assess our reconstruction method, we first evaluated the feasibility of inferring directed neural signaling from bulk transcriptomic data, because the “NeuronChat” algorithm uses single-cell-resolution datasets. We pseudo-bulked single-cell transcriptomic data from the mouse brain to predict the experimentally validated VISp and ALM connectomes, following the original study design. Our replication of the “NeuronChat” method, including pseudo-bulking and filtering for low-expression genes, yielded auROC = 0.8061 and auPRC = 0.8473 for the VISp model; auROC = 0.8333 and auPRC = 0.7913 for the ALM model (**Supplementary Fig. S4**). The original study reported auROC = 0.8316 and auPRC = 0.9153 for the VISp model; auROC = 0.7639 and auPRC = 0.7677 for the ALM model[31]. Pseudo-bulk estimation achieved predictive power comparable to the native single-cell implementation, indicating that ligand-receptor association-based inference retained performance at tissue-level resolution.

Based on these results, we benchmarked our proposed connectivity estimation method against tract-tracing-derived macaque connectome datasets, ‘Organized CoCoMac’ and ‘TheVirtualBrain’ (**Supplementary Fig. S5**). Detailed benchmark results and auxiliary measures are depicted in **Figs. 2–4** and **Supplementary Figs. S6, S7**. Our method’s performance exceeded that of the original “NeuronChat” algorithm on every point estimate for the Organized CoCoMac connectome (**Figs. 2A, 2B**), with auROC = 0.6868 (CI: [0.4763, 0.7300]) vs. 0.6426 and auPRC = 0.3308 (CI: [0.1595, 0.3882]) vs. 0.3010. All CIs are bootstrap 95% CIs (n = 1,000). The difference between the two methods became larger for TheVirtualBrain connectome (**Figs. 3A, 3B**), with auROC = 0.6842 (CI: [0.4563, 0.7375]) vs. 0.5212, and auPRC = 0.6953 (CI: [0.5123, 0.7635]) vs. 0.5195. Bootstrap intervals were wide; because both methods were applied to the same transcriptomes and benchmarked against the same reference connectomes, the observed improvement is attributable to our method rather than to the data.

**Fig. 2.**
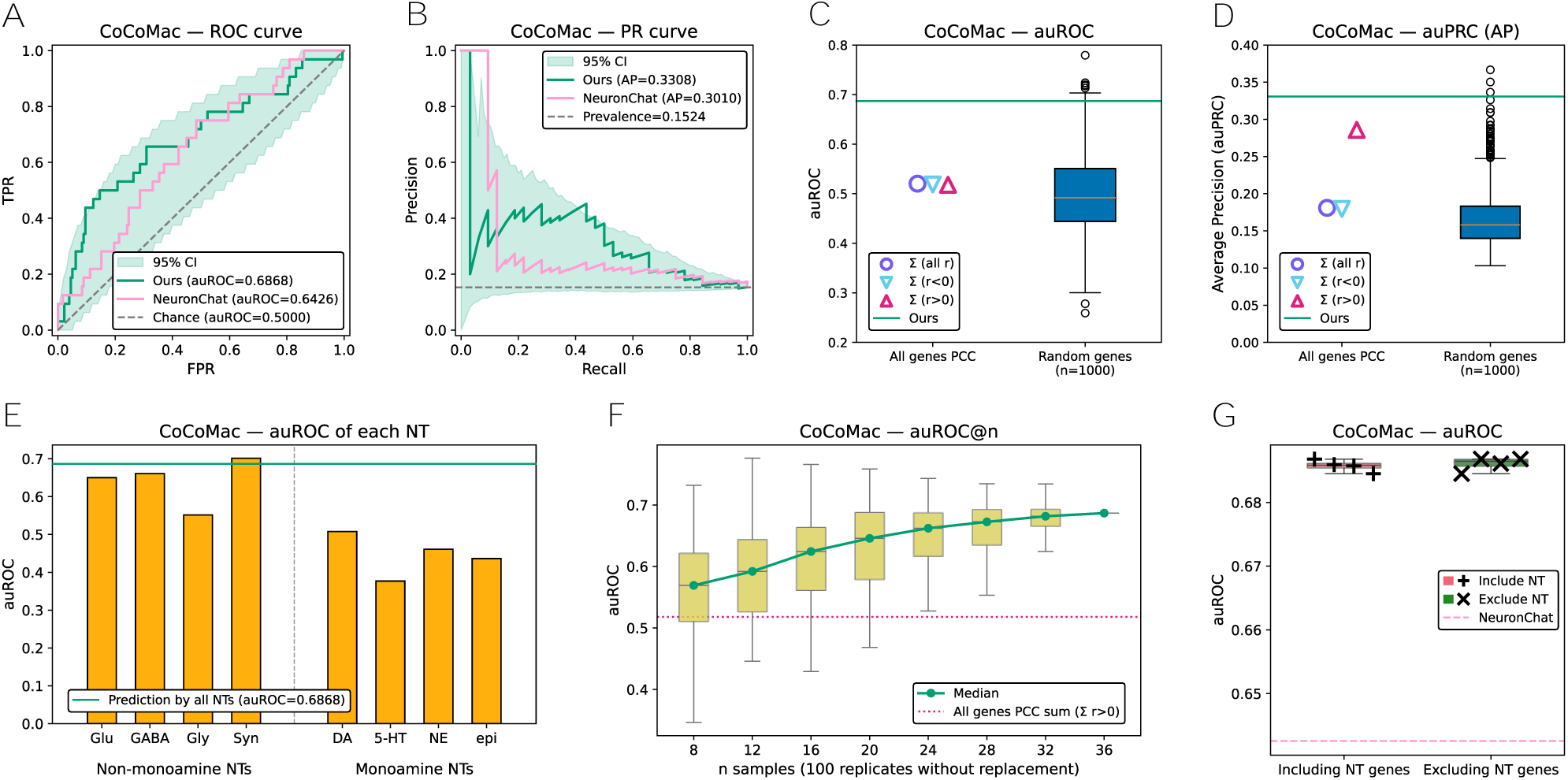
Benchmarking of connectivity estimates against the ‘Organized CoCoMac’ reference connectome. Connectivity strengths were estimated from macaque bulk RNA-seq data (GSE180092; 36 animals, 15 regions). Unless stated otherwise, strength estimates were summed across the eight neurotransmitter systems and synaptic adhesion for each region pair. Self-loops from subregion merging were excluded. (**A**) ROC and (**B**) PR curves for our pipeline and the “NeuronChat” method. Shading shows bootstrap 95% CIs for our pipeline (1,000 resamples). (**C**) auROC and (**D**) auPRC of three whole-transcriptome co-expression measures (left) and 1,000 random gene sets matched to the curated gene list (right). (**E**) auROC for each of seven neurotransmitter system and synaptic adhesion across 15 regions. Acetylcholine was excluded because the three regions with non-zero estimated abundance have only one positive reference edge. (**F**) auROC across subsample sizes (n = 8 to 36 animals in steps of 4; 100 subsamples drawn without replacement per size). The n = 36 point is the full cohort. (**G**) auROC across eight background gene panel sets (10 panels of 585 genes each), four drawn under each eligibility rule: genes expressed in both macaque and human data, with or without the 585 neurotransmission-related genes. auPRC counterparts of (E)-(G) are shown in Supplementary Fig. S6A-C.

**Fig. 3.**
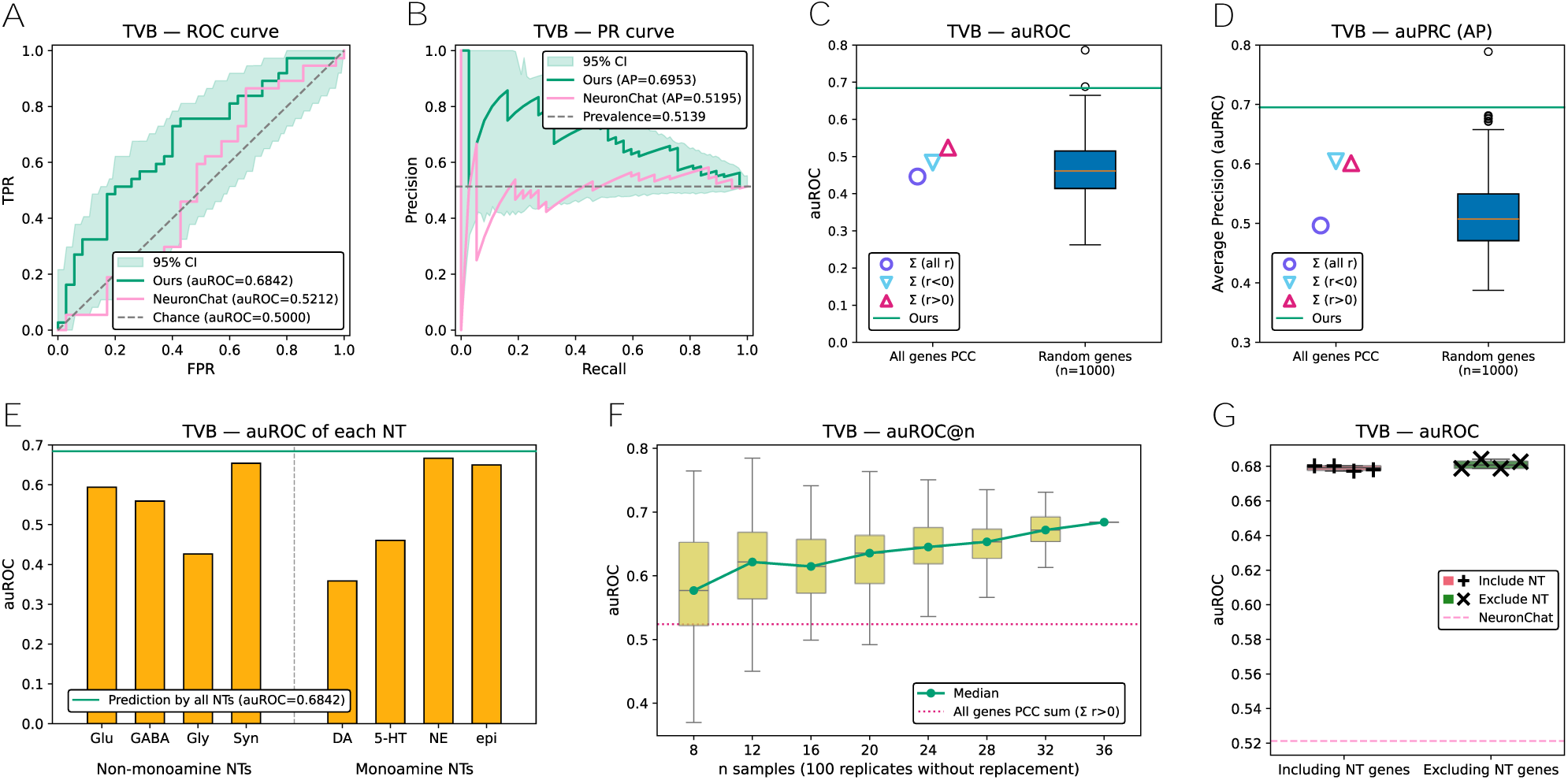
Benchmarking of connectivity estimates against ‘TheVirtualBrain’ reference connectome. Connectivity strengths were estimated from macaque bulk RNA-seq data (GSE180092; 36 animals, 15 regions). The estimates were compared against the right hemisphere portion of TheVirtualBrain connectome, matching the hemisphere sampled for transcriptomics. Unless stated otherwise, strength estimates were summed across the eight neurotransmitter systems and synaptic adhesion for each region pair. Self-loops from subregion merging were excluded. (**A**) ROC and (**B**) PR curves for our pipeline and the “NeuronChat” method. Shading shows bootstrap 95% CIs for our pipeline (1,000 resamples). (**C**) auROC and (**D**) auPRC of three whole-transcriptome co-expression measures (left) and 1,000 random gene sets matched to the curated gene list (right). (**E**) auROC for each of seven neurotransmitter system and synaptic adhesion across 15 regions. Acetylcholine was excluded because no reference edges are defined for the regions with non-zero estimated abundance. (**F**) auROC across subsample sizes (n = 8 to 36 animals in steps of 4; 100 subsamples drawn without replacement per size). The n = 36 point is the full cohort. (**G**) auROC across eight background gene panel sets (10 panels of 585 genes each), four drawn under each eligibility rule: genes expressed in both macaque and human data, with or without the 585 neurotransmission-related genes. auPRC counterparts of (E)-(G) are shown in Supplementary Fig. S7A-C.

**Fig. 4.**
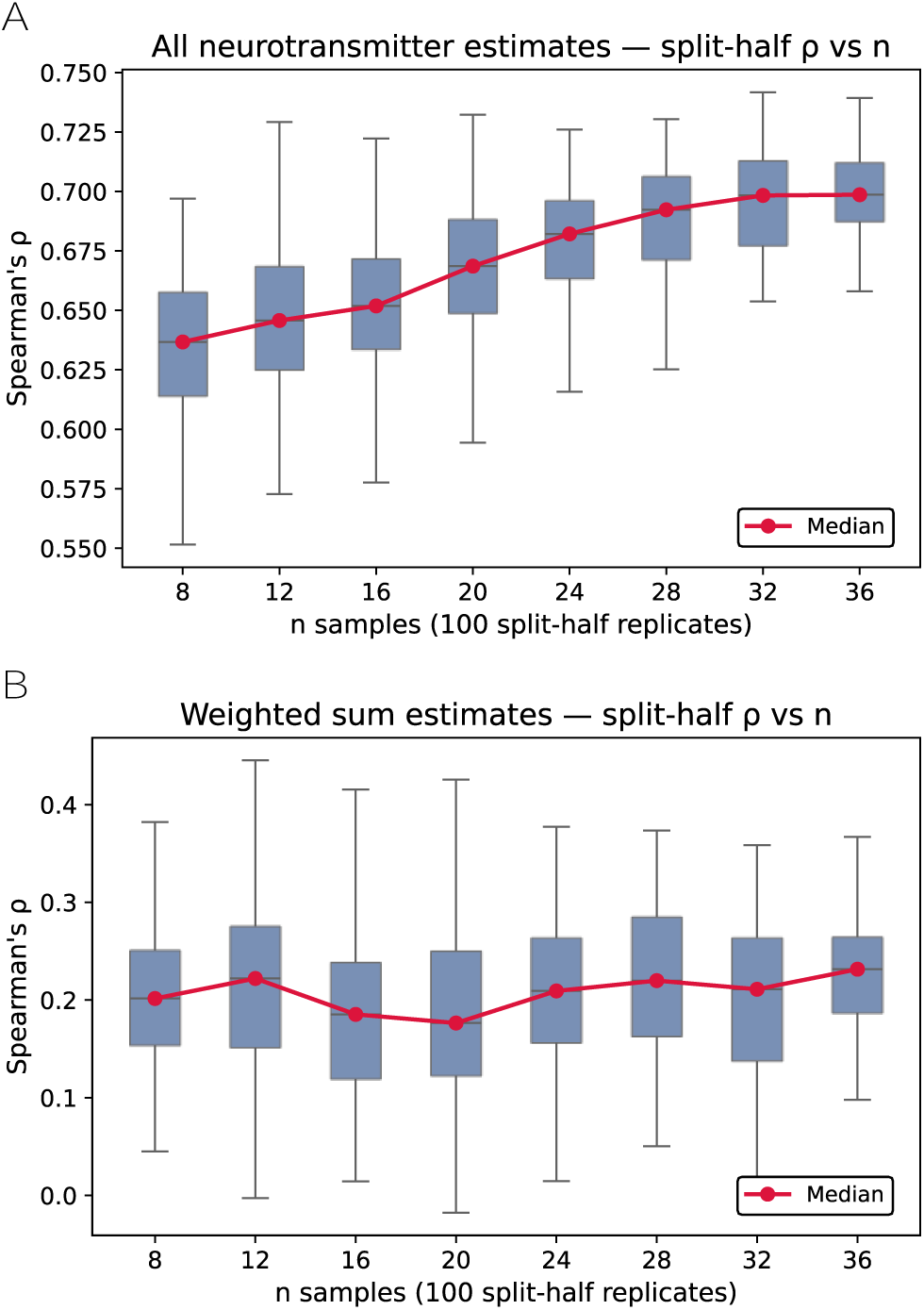
Agreement of macaque connectivity estimates between cohort halves across subsample sizes. Each of the 100 subsamples used in Figs. 2F and 3F (n = 8 to 36 animals in steps of 4) was split at random into two halves of n/2 animals. Connectivity strengths were estimated separately in each half, and agreement between the halves was quantified as Spearman’s ρ. At n = 36, the entire cohort was randomly split without subsampling. (**A**) Agreement across all neurotransmitter-by-region-pair connections. (**B**) Agreement of aggregated estimates in which the eight neurotransmitter systems and synaptic adhesion were summed for each region pair.

Our method also outperformed naïve interregional whole-transcriptome co-expression-based measures (**Figs. 2C, 2D, 3C, 3D, left side**). Three quantities were calculated: 1) sum of all PCCs for every possible gene pair, 2) sum of PCCs with r > 0, and 3) sum of PCCs with r < 0. The three quantities estimated interregional connectivity near the chance level (auROC: [0.4461, 0.5239]). The whole-transcriptome analysis indicates that interregional co-expression does not recover anatomical connectivity without a pre-defined gene list and background correction. Next, we randomly generated 1,000 gene sets and used them in estimating interregional connectivity strengths to calculate their predictive power (**Figs. 2C, 2D, 3C, 3D, right side**). Most of the random gene sets showed inferior performance compared to that of our curated neurotransmission-related gene list. Against any single reference connectome, no more than 14 randomly selected gene sets outperformed the curated gene list. None of those random gene sets outperformed the curated gene list when evaluated against the other reference connectome. Given that both connectomes were derived from the same tract-tracing database and their labels are positively correlated, this inconsistency indicates that the predictive performances of the random gene sets are not replicated. The incongruent performance reflects that the random gene sets are unstable.

We additionally analyzed the predictive performance of each neurotransmitter system and synaptic adhesion (**Figs. 2E, 3E, and Supplementary Figs. S6A, S7A**). Of the seven neurotransmitter systems and synaptic adhesion (acetylcholine excluded), four monoamine neurotransmitters showed lower predictive performance against the Organized CoCoMac connectome compared to non-monoamine neurotransmitters and synaptic adhesion (mean auROC: 0.4455 vs. 0.6410, mean auPRC: 0.1481 vs. 0.2842). Against TheVirtualBrain connectome, the gap narrowed for auROC and reversed for auPRC (mean auROC: 0.5339 vs. 0.5583; mean auPRC: 0.6912 vs. 0.5726). Monoamine neurotransmitters have limited sources across brain regions, whereas non-monoamine neurotransmitters have non-zero estimated abundance throughout regions, and neurexin expression is also widespread (**Supplementary Fig. S2**). To calibrate the source-region bias, we restricted the predicted connections and reference connectome to the 10 regions with a non-zero estimated abundance for every monoamine neurotransmitter and recalculated predictive performance (**Supplementary Figs. S6D, S6E**). In this additional analysis, the predictive power of non-monoamine neurotransmitters and synaptic adhesion remained almost unchanged (mean auROC = 0.6433), and the advantage of non-monoamine neurotransmitters and synaptic adhesion in predictive power was retained (mean auROC: 0.6433 vs. 0.4768). Each of the four non-monoamine auROC measures exceeded the highest monoamine auROC. The calibration analysis indicates that the low predictive power of each monoamine neurotransmitter does not originate from its source availability and is not an artifact of source and target region composition. When the same restriction was applied to TheVirtualBrain connectome, the dissociation was not reproduced (**Supplementary Figs. S7D, S7E**). Similar to the interpretation of the random gene set analysis, we interpret this disagreement as uncertainty between references arising from divergence between the two parcellations despite their shared source database.

We analyzed the sensitivity and robustness of our proposed method, as a small sample size could introduce subsampling variability and unstable correlation coefficients[80]. Accordingly, we designed a subsampling analysis and a half-split analysis for each subsample. As expected, predictive power (quantified as median auROC and auPRC) increased, and its variation (quantified as the interquartile range of the auROC and auPRC distributions) decreased as the number of samples increased. For the Organized CoCoMac connectome, median auROC already exceeded the whole-transcriptome co-expression baseline at n = 8, and median auPRC exceeded the baseline at n = 24 subsamples (**Fig. 2F and Supplementary Fig. S6B**). Results were also similar for TheVirtualBrain connectome (**Fig. 3F and Supplementary Fig. S7B**). For the half-split analysis, we calculated Spearman’s ρ using all possible neurotransmitter-by-region-pair connections, and using aggregated whole-neurotransmitter-level estimates. Connection-wise agreement increased as sample size increased, whereas aggregated estimates showed relatively low agreement between the two halves (**Figs. 4A and 4B**). To assess sensitivity to background genes, we repeated the full reconstruction pipeline and calculated benchmark performance across eight background gene panel sets. We drew four random panel sets under each of the two eligibility rules, which include and exclude neurotransmission-related genes (**Figs. 2G, 3G, and Supplementary Figs. S6C, S7C**). Benchmark performance was stable across all random draws and exceeded that of the “NeuronChat” algorithm in every case. Deviations across panels were small in predictive power metrics (standard deviation < 0.0072); the coefficient of variation (CV) was also small (CV < 2.18%) for all four metrics. The two candidate background gene eligibility rules gave almost identical results (mean auROC difference < 0.003), and estimated connectivity strengths were highly concordant across all panels (pairwise PCC > 0.9993). The low performance variation across background panels suggests that the estimates were not highly sensitive to the random draw and candidate pool generation. The benchmarking analysis showed that transcriptomic connectivity estimates were concordant with experimentally validated anatomical connections, and remained stable across subsamples and background panels. We applied this method to a human cohort to map the “interregional molecular connectome” and analyze the dysregulations in patients with schizophrenia.

### Interregional neurotransmitter signaling dysregulations in schizophrenia

We next applied this method to a large-scale human transcriptomic cohort (1,037 donors, 628 with region-paired samples; **Table 1 and Supplementary Table S1**) to map connectivity changes across the sampled regions in schizophrenia. We quantified the weighted connectivity difference of each connection between diagnostic groups and assessed its statistical significance. Empirical null distributions for each connection were approximately symmetric (skewness: [−0.5408, +0.3782]) and platykurtic (mean kurtosis: −0.3463). All metrics, including raw and standardized connectivity differences, their label-permutation quantiles, permutation p-values, BH-adjusted q-values, and abundance estimate statistics, are provided in **Supplementary Table S4**. Several neurotransmitters have zero estimated abundance in some regions, which limits the number of connections that could be tested among the 180 total possible connections ((8 neurotransmitter systems plus synaptic adhesion) × 5 source regions × 4 target regions). All the sampled regions lacked expression of the core synthetic enzymes for epinephrine and norepinephrine, yielding zero abundance estimates. Estimated abundance was also zero for dopamine, serotonin, and acetylcholine in the cortical regions, for serotonin in CN, and for acetylcholine in HP. In each case, one or more core synthetic enzyme genes fell below the expression threshold. Therefore, a zero estimate reflects filtered expression and does not necessarily mean that the neurotransmitter is absent from the region. These zero estimates excluded 72 connections (20 each for adrenergic and noradrenergic, 12 each for serotonergic and cholinergic, and 8 dopaminergic), yielding a subset of 108 testable connections.

Of the 108 testable connections, 12 connections reached the nominal significance threshold. We then applied the directional consistency filter that retains connectivity differences consistent with neurotransmitter abundance differences. A total of 10 connections remained after applying the filter (**Fig. 5A**). The 10 connections were considered candidate dysregulations in patients with schizophrenia (**Figs. 5B–5D**). The filter excluded two candidates whose direction of change in abundance estimates was opposite to the weighted connectivity difference. The two excluded candidates were an increased GABAergic CC → HP connection and a reduced glycinergic PFC → CC connection. For these two connections, the group difference in estimated abundance of the corresponding neurotransmitter systems contradicts the direction of connection differences (Hedges’ g = −0.3428 and +0.4372, respectively, both t-test p < 0.05). The directional consistency filter was applied solely based on statistical significance: the glutamate abundance estimate in NAc was reduced with a low-to-modest effect size (Hedges’ g = −0.3066, t-test p = 0.1084), but the elevated glutamatergic NAc → CC connection was retained. The filter therefore behaved as a specificity check on directional consistency. However, two of the 10 candidates (glycinergic CN → HP and CC → CN synaptic adhesion) had two-tailed p-values between 0.05 and 0.1. We interpret these 10 candidates as exploratory, data-driven findings for hypothesis generation.

**Fig. 5.**
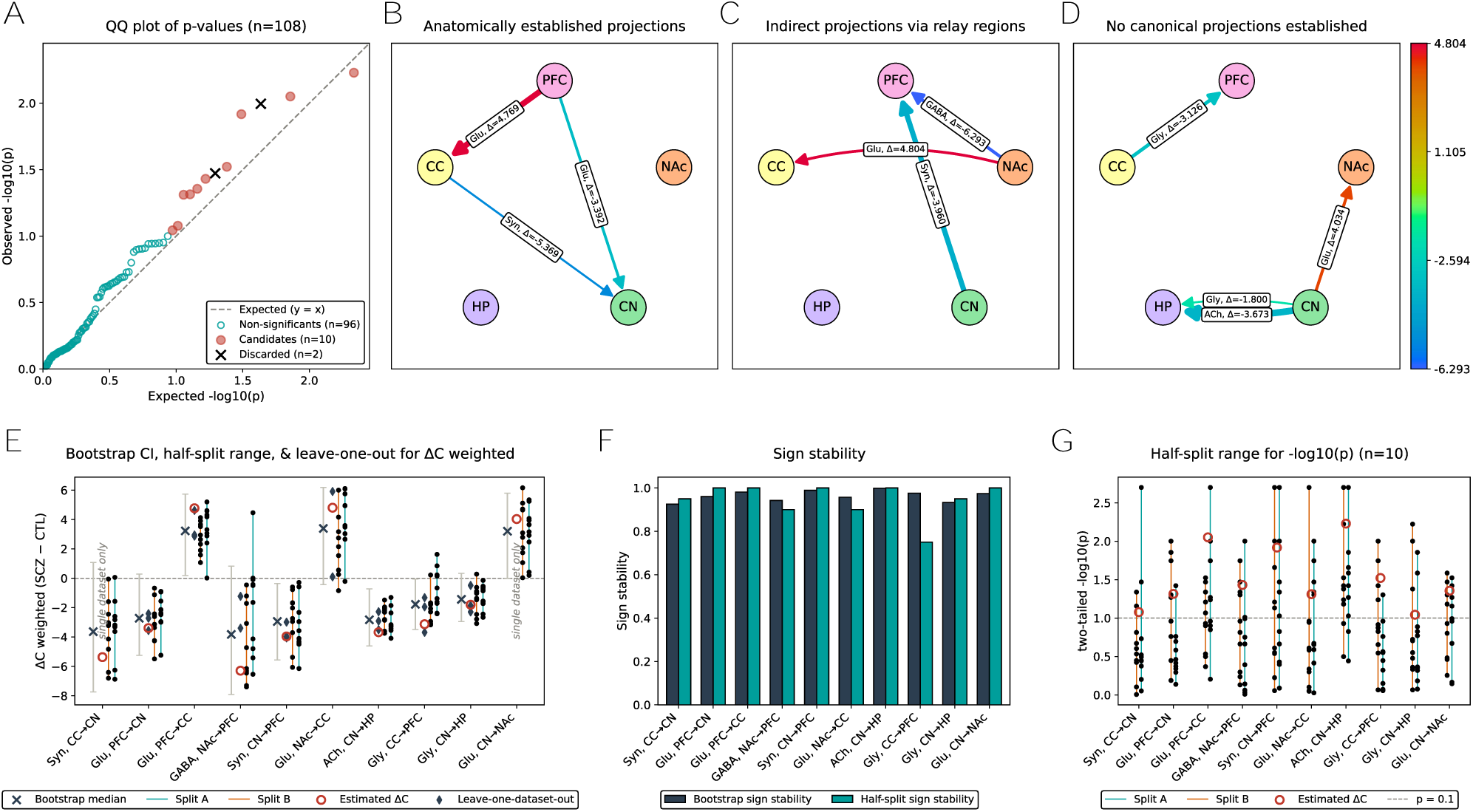
Candidate interregional connectivity dysregulations in schizophrenia and robustness of their difference scores. The difference score is the weighted strength estimate of controls subtracted from that of patients with schizophrenia. (**A**) Quantile-quantile plot of two-tailed permutation p-values for the 108 testable connections (10,000 diagnostic label permutations). Of these, 12 reached significance threshold (label-permutation quantile < 0.05 or > 0.95); 10 passed the directional consistency filter and became candidates; 2 were excluded by the filter. (**B-D**) Candidates grouped by anatomical support. (**B**) Three candidates aligned with established anatomical connections. (**C**) Three candidates aligned with indirect connections through relay regions. (**D**) Four candidates lacked an established monosynaptic connection. Edge color represents the magnitude of the difference score, and edge width represents significance from label permutation analysis. (**E**) Difference scores of each candidate in the full cohort, with bootstrap 95% CIs (1,000 resamples), leave-one-dataset-out estimates, and half-split estimates from 10 splits. Leave-one-dataset-out estimates are shown for the eight candidates with two or more contributing datasets. The two that rest on a single dataset are marked “single dataset only”. (**F**) Sign stability, the proportion of the resamples retaining the sign of the full-cohort estimates, for bootstrap resampling (1,000 resamples) and the 20 halves. (**G**) − log10(p) of the two-tailed permutation p-value of each candidate in each of the 20 halves (500 label permutations per half).

As in the macaque benchmarking analysis, we additionally analyzed the robustness and sensitivity of our findings. Three subsampling analyses tested whether the observations were stable under sample exclusion: leave-one-dataset-out, bootstrap resampling (n = 1,000), and half-split (**Figs. 5E–5G**). When one dataset was omitted and difference scores were recalculated, the eight candidates with two or more contributing datasets retained their difference sign on each exclusion (**Fig. 5E**). The other two rest on a single dataset. For the bootstrap resampling, the signs of difference scores remained largely unchanged (**Fig. 5E**), with sign stability (defined as the proportion of the n = 1,000 resamples retaining the sign of the point estimate) exceeding 92.5% for every candidate (**Fig. 5F**). Only four of the 10 candidate connections (glutamatergic PFC → CC, cholinergic CN → HP, glycinergic CC → PFC, and CN → PFC synaptic adhesion) had 95% CIs that excluded zero. The sign of the difference is therefore stable for all 10 candidates, whereas the magnitude is distinguishable from zero for only four. In the half-split analysis (**Fig. 5F**), the consistency declined (sign stability for the 20 halves > 75%). These sample exclusion analyses show that the 10 candidate connections are not single-dataset-driven artifacts and are not heavily dependent on cohort composition. Although directions of the difference scores were stable throughout the subsampling analyses, their nominal significance was more fragile: the 10 connections had two-tailed p-values < 0.1 for 20%–75% of the 20 halves (**Fig. 5G**). Finally, we repeated the full human analysis (10,000 label permutations) under two candidate background gene eligibility rules (including and excluding neurotransmission-related genes, see Methods for details) to assess sensitivity to the rules. All 10 candidate dysregulations met the nominal significance threshold in the two configurations. Connection-level difference scores were almost identical across the two configurations (PCC = 0.9995). The concordance between the two rules matches findings from the macaque connectome benchmarking analysis, and indicates that the schizophrenia-specific dysregulations were not highly sensitive to the candidate background gene eligibility rules. These auxiliary sensitivity analyses showed that the signs of the candidate difference scores were stable under sample restriction, dataset exclusion, and candidate background gene eligibility rules.

To assess the stability of our difference score calculation, we computed Spearman’s ρ between half splits using all the testable connections for each diagnosis group and for the difference scores. Across 108 connections for 10 splits, mean ρ was 0.6184 for healthy controls, 0.5842 for patients with schizophrenia, and 0.0927 for the differences between two diagnostic groups. A Spearman’s ρ of ∼0.1 is low in absolute terms, so individual difference scores are not stable at this sample size. However, it exceeded 99.3% of the values in the empirical null distribution (permutation n = 500) obtained from the diagnostic label shuffle. The high empirical percentile of the observed ρ indicates that the overall pattern of differences was replicated between independent halves. We next partitioned the 108 connections by the number of contributing datasets: three or more (samples ≥ 133; 40 connections), two (83 ≤ samples ≤ 109; 32 connections), and one (samples ≤ 65; 36 connections), and report ρ within each partition. For the first two groups, half-split agreement (mean ρ) was comparable to that from 108 connections, and consistently larger than the null (mean ρ: 0.1055 and 0.1239, quantiles: 96.91% and 97.31%, respectively). For the third group, agreement was near-zero and indistinguishable from the null distribution (mean ρ = −0.0147, quantile = 40.62%).

We note that these measures are not directly comparable between the two species. In the macaque benchmarking (**Figs. 2F, 4A**), half-split analysis resulted in a median Spearman’s ρ of 0.6821 at n = 24, with a median auROC of 0.5920 at n = 12 (i.e., each half of a 24-animal subsample). The macaque half-split agreement was only calculated from connectivity estimate scores, whereas the human half-split agreement was calculated from difference scores that compound the sampling variability. Consistent with this, the human diagnosis-group-level estimates reached Spearman’s ρ of 0.6184 and 0.5842, which are comparable to the macaque value. The reduction to 0.0927 arises only when the two group-level estimates are differenced. The macaque value is therefore interpreted as an upper bound on what our method can achieve at a comparable number of samples, not as an equivalent quantity. In the macaque benchmarking, high agreement between halves coexisted with modest anatomical accuracy. This coexistence shows that half-split agreement indicates stability and says little about validity against the reference connectome. The low human half-split agreement is also expected from sample size. Given that the region pair with the lowest number of samples comprised 25 controls and 29 patients with schizophrenia (halved down to 12 donors in half-split analysis), we do not read the low Spearman’s ρ as a failure to reproduce the underlying signal.

Of the 10 candidate dysregulations, which span four neurotransmitter systems and synaptic adhesion, seven showed decreased and three showed increased connectivity strength estimates in patients with schizophrenia (**Figs. 5B–5D**). Epinephrine and norepinephrine were excluded from the connectivity difference analysis pipeline because the sampled regions lacked the expression of core synthetic enzymes, yielding zero abundance estimates. No dopaminergic or serotonergic connection reached nominal significance based on our permutation analysis and directional consistency filter. Most of our candidates show bidirectional dysregulation in cortico-striatal connections. This pattern is consistent with involvement of the CSTC circuit in schizophrenia pathophysiology. The cortico-striatal concentration of our candidate dysregulations (8 of the 10 candidates) partly reflects the composition of the sampled regions: four of the five are cortical or striatal, and 63 of the 108 testable connections lie among them. The majority of these dysregulations were driven by changes in co-expression coordination, as reflected by the low-to-modest effect sizes for group differences in neurotransmitter abundance estimates (|Hedges’ g| ≤ 0.3108). We then stratified those candidate dysregulations by the degree of established evidence. The first group comprises three connections: an increased intra-cortical glutamatergic connection, a decreased cortical glutamatergic output to CN, and a decreased synaptic adhesion in cortical output to CN, all supported by previous findings. The second group includes ascending striatal outputs to cortex, which are mediated through relay regions such as the thalamic nuclei, the pallidum, and the midbrain. The final group encompasses an increased intra-striatal glutamatergic connection and reduced cholinergic and glycinergic connections. None of these are supported by established monosynaptic pathways (see Discussion).

### LLM-augmented screening quantifies literature support for biomedical statements

We tested whether the selected model “gpt-oss-120b” could classify an abstract as supporting, refuting, or neutral towards a given hypothesis, and whether the literature screening framework could quantify the degree of evidence consensus for the well-established biomedical hypotheses against the literature corpus. We first benchmarked the NLI capability of the selected model on a biomedical-specific NLI benchmark dataset named BioNLI[75]. This dataset is designed to evaluate an LLM’s ability to distinguish true biological claims from subtle semantic inversions and from unrelated sentences. The discrimination ability is critical for evidence synthesis in the biomedical domain. The selected model achieved an accuracy of 0.8972 (**Fig. 6**). The accuracy fell between that of the weakest-performing model “deepseek-v3.2-thinking” (accuracy = 0.8500) and the best-performing model “gemini-3-pro-preview-11-2025-high” (accuracy = 0.9450) of the 23 selected models evaluated in the ‘Believe’ framework[37].

**Fig. 6.**
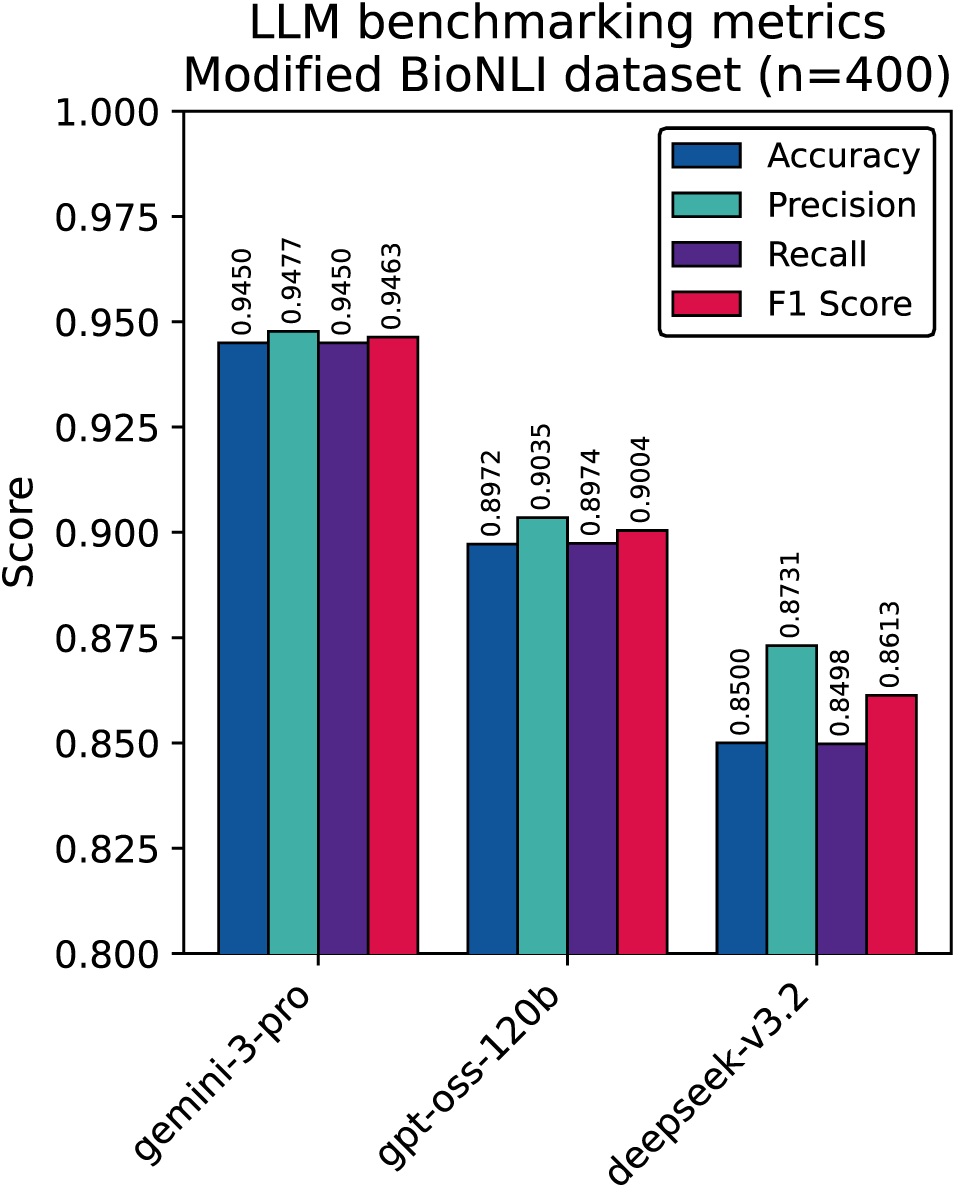
Natural language inference performance of three LLMs on the BioNLI dataset. Accuracy, macro precision, macro recall, and macro F1 score for three LLMs: “gpt-oss-120b”, the model used for literature screening; “gemini-3-pro-preview-11-2025-high”, the best-performing model; and “deepseek-v3.2-thinking”, the weakest-performing model among the 23 models evaluated in the ‘Believe’ framework. Each model classified whether an abstract supports, refutes, or is neutral towards a paired hypothesis in the modified BioNLI dataset (400 instances: 134 positive, 133 negative, and 133 neutral).

Next, we assessed two proposed calibration metrics: Alignment, representing the strength of consensus for a given hypothesis, and Relevancy, representing the volume of relevant literature. We observed consistent patterns across all calibration hypotheses regardless of literature volume (**Table 2**). For example, the canonical biomedical observation that “Patients with Type 2 Diabetes Mellitus exhibit insulin resistance” yielded an Alignment score of 0.9935 and a Relevancy score of 0.0556, indicating that supporting abstracts strongly dominated the relevant literature corpus. The near-perfect Alignment score indicates that the screening recovered the established consensus for this hypothesis. The adversarial negation of the premise that “Patients with Type 2 Diabetes Mellitus do not exhibit insulin resistance” resulted in an Alignment score of 0.0056 and a Relevancy score of 0.0665, indicating that our framework correctly identified this hypothesis as contradicted by strong consensus. Similarly, another negation, “Patients with Type 2 Diabetes Mellitus exhibit high insulin sensitivity,” resulted in an Alignment score of 0.0033 and a Relevancy score of 0.0540, indicating that the framework is capable of identifying lexical negation regardless of how the negation is represented in natural language. The calibration examples produced the expected direction of Alignment scores independent of literature volume, supporting use of these metrics as indicators for quantifying literature support for our findings. The benchmarking results indicate that the selected model approaches the performance of the best-performing proprietary model, and that it assigns the expected direction of Alignment to canonical hypotheses and their negations. These properties support the use of LLM-augmented literature screening as a systematic, scalable approach to evidence synthesis.

**Table 2.**
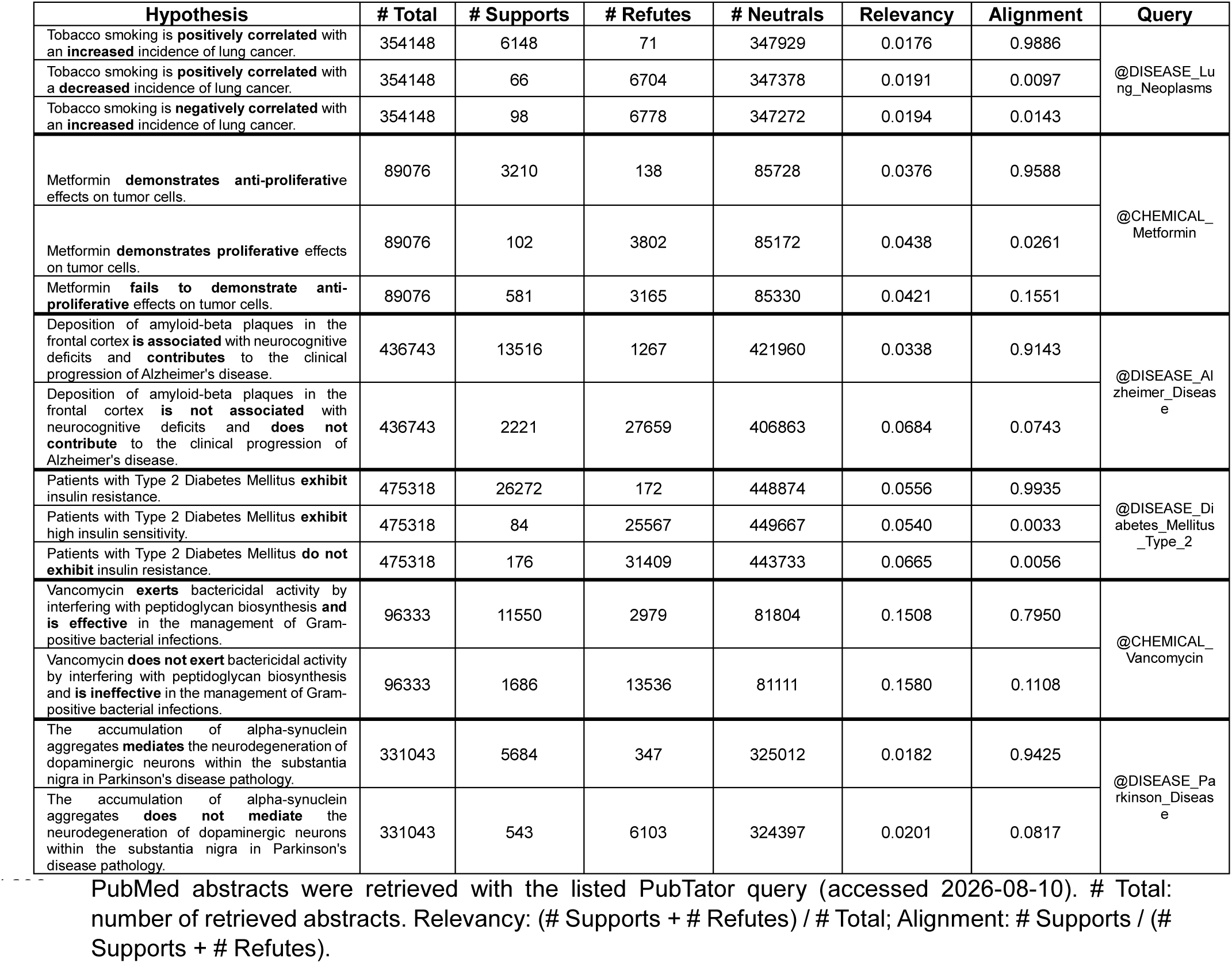
LLM literature screening of well-established biomedical hypotheses and their adversarial negations.

Applying the LLM-augmented literature screening framework to our 10 candidates, we calculated the degree of literature support for our observations using the proposed calibration metrics (**Table 3**). To systematically synthesize evidence for the findings, we classified the candidates into three distinct categories based on calibration metrics. The first category encompasses literature-supported hypotheses for which known literature evidence supports our candidates, characterized by moderate-to-high Alignment scores (Alignment ≥ 0.5) and Relevancy > 0.0001. The first category includes three decreased candidates: glutamatergic PFC → CN, GABAergic NAc → PFC, and cholinergic CN → HP. This category shows agreement between our observations and previously reported abnormalities from post-mortem and neurochemical studies, while adding specificity to circuit-level mechanisms. The second category includes controversial mechanisms, characterized by low Alignment scores (Alignment < 0.5) and Relevancy > 0.0001. Three increased glutamatergic dysregulations fall here: intra-cortical PFC → CC, intra-striatal CN → NAc, and striatal output NAc → CC. These low Alignment scores suggest either disagreement about the directed dysregulations or their direct refutation in existing literature. The third category comprises underexplored predictions with little directly relevant literature, with very low Relevancy (Relevancy < 0.0001). The third category includes two glycinergic connections (CN → HP and CC → PFC), and two synaptic adhesion dysregulations: one within the cortico-striatal circuit (CC → CN) and one in striatal output to cortex (CN → PFC). These candidates point to mechanisms that require experimental validation. We also note that the amount of relevant and supporting literature is not directly concordant with anatomically established connections. Well-studied neurotransmitters (glutamate, GABA, and acetylcholine) have relatively more relevant articles compared to glycine and synaptic adhesion, although the synaptic adhesion candidates align with direct or indirect anatomical connections documented by several independent lines of evidence. Our literature screening quantifies the amount of evidence available for each candidate and the direction in which that evidence points. This quantification does not establish or rank our candidates, and does not provide anatomical evidence for the findings.

**Table 3.** LLM literature screening of the 10 candidate dysregulations in schizophrenia.

| Hypothesis | # Supports | # Refutes | # Neutrals | Relevancy | Alignment |
| --- | --- | --- | --- | --- | --- |
| Glutamatergic signaling from the prefrontal cortex to the caudate nucleus is hypofunctional in patients with schizophrenia. | 156 | 76 | 245842 | 0.000943 | 0.6724 |
| GABAergic signaling from the nucleus accumbens to the prefrontal cortex is hypofunctional in patients with schizophrenia. | 23 | 18 | 246033 | 0.000167 | 0.5610 |
| Cholinergic signaling from the caudate nucleus to the hippocampus is hypofunctional in patients with schizophrenia. | 27 | 18 | 246029 | 0.000183 | 0.6000 |
| Glutamatergic signaling from the prefrontal cortex to the cingulate cortex is hyperfunctional in patients with schizophrenia. | 62 | 862 | 245150 | 0.003755 | 0.0671 |
| Glutamatergic signaling from the caudate nucleus to the nucleus accumbens is hyperfunctional in patients with schizophrenia. | 14 | 279 | 245781 | 0.001191 | 0.0478 |
| Glutamatergic signaling from the nucleus accumbens to the cingulate cortex is hyperfunctional in patients with schizophrenia. | 13 | 303 | 245758 | 0.001284 | 0.0411 |
| Glycinergic signaling from the cingulate cortex to the prefrontal cortex is hypofunctional in patients with schizophrenia. | 7 | 3 | 246064 | 0.000041 | 0.7000 |
| Glycinergic signaling from the caudate nucleus to the hippocampus is hypofunctional in patients with schizophrenia. | 2 | 1 | 246071 | 0.000012 | 0.6667 |
| Synaptic adhesion integrity in the projections from the cingulate cortex to the caudate nucleus is decreased in patients with schizophrenia. | 7 | 1 | 246066 | 0.000033 | 0.8750 |
| Synaptic adhesion integrity in the projections from the caudate nucleus to the prefrontal cortex is decreased in patients with schizophrenia. | 17 | 1 | 246056 | 0.000073 | 0.9444 |

### Bidirectional dysregulation of the CSTC circuit in schizophrenia

Six of the 10 candidate dysregulations are supported by well-established anatomical connections or connections mediated through relay regions. The six dysregulations together span the CSTC circuit (**Figs. 5B, 5C**), which is consistently implicated in schizophrenia[81]. Of these dysregulations, three had well-established anatomical evidence, whereas the remaining three were within striato-cortical connections through relay regions. These were not uniform in direction: cortical and striatal output to CC were increased, but connections targeting CN or PFC were decreased. First, increased intra-cortical glutamatergic connectivity and reduced cortico-striatal connections were identified (**Fig. 5B**). These connections were supported by well-established anatomical evidence. The glutamatergic intra-cortical PFC → CC connection strength was increased in patients with schizophrenia (Δ*C^W^* = +4.7690, quantile = 0.9956). In contrast, the glutamatergic cortical output PFC → CN connection strength was decreased (Δ*C^W^* = −3.3917, quantile = 0.0242), as was synaptic adhesion strength in the CC → CN connection (Δ*C^W^* = −5.3694, quantile = 0.0418). Group differences in neurotransmitter abundance estimates were not statistically significant and were of negligible effect size in the three connections (|Hedges’ g| ≤ 0.1095, all t-test p > 0.1045), indicating that the difference in connectivity estimates was predominantly driven by changes in co-expression coordination, not by changes in synthesis capacity.

Next, dysregulations in ascending striatal outputs to the cortex aligned with the known CSTC circuit, which connects these regions through relay regions, including the pallidum, the midbrain ventral tegmental area (VTA), and thalamic nuclei (**Fig. 5C**). These findings highlight changes in co-expression coordination along an indirect ascending pathway from striatal regions to cortical regions. The glutamatergic NAc → CC connection strength was increased (Δ*C^W^* = +4.8043, quantile = 0.9756). In contrast, the GABAergic NAc → PFC connection strength was decreased (Δ*C^W^* = −6.2931, quantile = 0.0185), as was synaptic adhesion strength in the CN → PFC connection (Δ*C^W^* = −3.9597, quantile = 0.0060). As with the cortical inputs to striatal regions, neurotransmitter abundance estimates remained nearly unchanged for decreased connections (|Hedges’ g| ≤ 0.1447, all t-test p > 0.4296), but the glutamate abundance estimate in NAc was reduced with a low-to-modest effect size (Hedges’ g = −0.3066, t-test p = 0.1084). Because the abundance difference did not reach the nominal significance threshold, the connection was retained. The direction of this abundance estimate is opposite to the direction of co-expression coordination increase, suggesting that this candidate is the least internally consistent of the 10 candidates.

We prioritized genes that contribute to the changes in co-expression coordination. The reduced glutamatergic PFC → CN connection was predominantly driven by the presynaptic vesicular transporter SLC17A7 (VGLUT1) with postsynaptic metabotropic receptors (GRM4 and GRM5) and ionotropic receptors (α-amino-3-hydroxy-5-methyl-4-isoxazolepropionic acid (AMPA) receptor GRIA2 and N-methyl-D-aspartate (NMDA) receptor GRIN1). The two elevated glutamatergic connections were also driven by SLC17A7, accompanied by SLC17A6 (VGLUT2) and the synthesis enzyme GLS. Postsynaptic prioritized genes of the elevated glutamatergic connections in CC spanned multiple receptor classes (metabotropic receptors GRM3, GRM4, GRM8, kainate receptors GRIK3, GRIK4, GRIK5, and NMDA receptor GRIN2A). The prioritized genes were statistically enriched in related glutamate receptor subclass GO terms (GO:0015277, “kainate selective glutamate receptor activity” for elevated connections targeting CC; GO:0007196, “adenylate cyclase-inhibiting G protein-coupled glutamate receptor signaling pathway” for the decreased connection targeting CN). The prioritized presynaptic genes were enriched in glutamate transporter activity and glutaminase-related GO terms (GO:0140788, “L-glutamate uniporter activity” and GO:0004359, “glutaminase activity”). For the GABAergic output to PFC, SLC32A1 (VGAT) drove the dysregulation alongside system-specific postsynaptic receptor subunits (GABA-A receptor subunits GABRA3, GABRB1, GABRB3, GABRG1, and GABA-B receptor subunit GABBR1). For the synaptic adhesion deficits, different neurexins and neuroligins (neurexins NRXN2 and NRXN3; neuroligins NLGN2, NLGN3, and NLGN4X) predominantly drove the reduction in these connections. The difference in prioritized contributing genes between the cortical-to-striatal connections (NRXN3) and striatal-to-cortical connections (NRXN2) in synaptic adhesion dysregulations suggests divergent roles for each neurexin gene in region-specific synapse specification. The prioritized genes were not significantly enriched in any GO term for the GABAergic connection or synaptic adhesion deficits.

The LLM screening indicated mixed literature consensus for these candidates, regardless of whether corresponding anatomical connections were directly or indirectly established (**Table 3**). Although the decreased glutamatergic PFC → CN connection showed an Alignment score of 0.6724 (156 supporting, 76 refuting articles), two increased glutamatergic connections (PFC → CC connection and NAc → CC connection) each had a low Alignment score (0.0671 and 0.0411, respectively). The discrepancy in Alignment scores suggests that increased glutamatergic connectivity captured at the transcriptomic level reflects a more complex mechanism than a simple downstream consequence of NMDA receptor hypofunction. These candidates need further investigation. The literature screening showed that biomedical consensus moderately supports the decreased GABAergic NAc → PFC connection (Alignment score of 0.5610, with 23 supporting and 18 refuting). Synaptic adhesion deficits in the CC → CN and CN → PFC connections each returned a low Relevancy score (Relevancy < 0.0001). The low Relevancy indicates that although CSTC circuit dysfunction and synaptic loss have been theoretically implicated in schizophrenia, specific changes in interregional connectivity remain largely unexplored despite the anatomical evidence for these connections.

### Altered co-expression coordination without an established monosynaptic connection

The remaining four out of 10 candidates were not explicitly supported by established monosynaptic connections between source and target regions (**Fig. 5D**). These included an intra-striatal glutamatergic coordination increase between CN and NAc (Δ*C^W^* = +4.0339, quantile = 0.9780), along with a cholinergic coordination decrease between CN and HP (Δ*C^W^* = −3.6733, quantile = 0.0029). The other two connections were decreased glycinergic coordination, one between CC and PFC and the other between CN and HP (Δ*C^W^* = −3.1257 and −1.7995, quantile = 0.0150 and 0.0452, respectively). Group differences in neurotransmitter abundance estimates were not significant for the three connections involving CN (|Hedges’ g| ≤ 0.1574, all t-test p > 0.0860), although glycine abundance estimates in CC were reduced with a low-to-modest effect size (Hedges’ g = −0.3108, t-test p = 0.0738). The non-significant group differences in abundance estimates across these connections indicate that the co-expression coordination changes were not primarily driven by changes in synthesis capacity in the source region. Several transporter genes and synthesis enzymes contribute to the dysregulation (glutamate: SLC17A7 (VGLUT1); acetylcholine: SLC18A3 (VAChT) and ACHE; glycine: SLC6A9 (GlyT1), SLC32A1 (VGAT), DLD, SHMT2). Distinct system-specific postsynaptic receptor subunits were prioritized (glutamate: ionotropic NMDA receptor subunits GRIN2D, GRIN3A, AMPA receptor subunit GRIA1, metabotropic receptor GRM7; acetylcholine: nicotinic receptor subunits CHRNA3, CHRNA5, CHRNA7, CHRNB2; glycine: GLRB). Prioritized genes of increased glutamatergic coordination between CN and NAc were statistically enriched in GO terms that are related to glycine (GO:0070905, “serine binding”, GO:0160212, “glycine-gated cation channel activity”, and GO:0017146, “NMDA selective glutamate receptor complex”). These enriched terms suggest that the intra-striatal glutamatergic and glycinergic changes are molecularly linked, although the glycinergic coordination between CN and NAc was not considered a candidate dysregulation. For other connections, prioritized genes were not significantly enriched in any GO term.

LLM screening showed that these coordination changes were not uniformly supported by literature evidence (**Table 3**). Glutamatergic and cholinergic coordination (between CN and NAc, and between CN and HP) both resulted in a sufficient number of relevant articles (Relevancy > 0.0001). Glutamatergic co-expression changes were poorly supported (Alignment = 0.0478, with 14 supporting and 279 refuting), whereas cholinergic changes were moderately supported (Alignment = 0.6000, with 27 supporting and 18 refuting). Glycinergic coordination changes each returned a low Relevancy score (Relevancy < 0.0001). Because our source-to-target labels are based on the functional classification of presynaptic and postsynaptic neurotransmission-related genes, all four candidates in this section should be interpreted as coordinated transcriptional regulation of presynaptic and postsynaptic machinery between regions, not as direct monosynaptic projections.

## Discussion

Our analysis of large-scale transcriptomic data from 1,037 individuals is consistent with a pattern of altered interregional co-expression coordination that maps onto the CSTC circuit. We first benchmarked our method using transcriptome datasets paired with two macaque connectomes derived from tract-tracing studies. Our method predicted anatomical connectivity more accurately than measures based on either the whole transcriptome or randomly selected gene sets. We then identified 10 candidate dysregulated interregional neurotransmission connections, six of which align with established direct or indirect connections. These connections describe a bidirectional pattern: co-expression coordination of neurotransmission-related genes on inputs to CC was increased, whereas cortical output to CN and striatal output to PFC were reduced. The observed circuit-level dysregulations are compatible with the CSTC circuit hypothesis and may help identify distinct molecular and anatomical mechanisms underlying schizophrenia pathophysiology. This pattern also supports the perspective of schizophrenia as a multi-regional, multi-neurotransmitter disorder with disconnected neural circuits. We then used an LLM-augmented literature screening framework to quantify the amount and direction of supporting evidence for each candidate connection. The framework calibrated the degree of existing evidence for each candidate. The calibration showed that some of our observations are supported by established findings, while others point to mechanisms with little directly relevant literature. We present these findings as exploratory and hypothesis-generating[82], and interpret them below primarily through their anatomical coherence, contributing-gene composition, and concordance with independent literature, along with our literature screening results.

The most noticeable feature of our candidates is their coherence with the CSTC circuit. The circuit is implicated in schizophrenia through the loss of functional connectivity. The CSTC circuit spans several interregional connections spanning cortical, thalamic, hippocampal, striatal, midbrain, and pallidal regions, and the amygdala[81]. The circuit affects the midbrain dopaminergic neurons, and then reciprocally influences striatal and cortical regions[81]. Among our observations, reduced glutamatergic PFC → CN connectivity corresponds to one of the most reproducible circuit-level findings in schizophrenia neuroimaging. Reduced functional and structural connectivity between the PFC and the dorsal associative striatum has been repeatedly demonstrated, and its normalization tracks response to antipsychotic treatment[83–85]. The agreement between neuroimaging studies and our finding for the same connection provides the clearest external support among our candidates. This reduction also coexists with increased excitatory coordination towards CC. This pattern might arise when descending cortical output to the striatum is weakened but excitatory striatal coordination is preserved or even increased. Elevated glutamatergic metabolite levels in the basal ganglia have been reported[24, 86], although metabolite concentrations and co-expression estimates are not directly comparable. Decreased striatal GABAergic output to cortex could participate in this reciprocal loop, a possibility supported by molecular observations including altered GABAergic interneuron markers[87] and reduced expression of GABA synthesis and transport genes[88, 89]. Cortico-striatal dysfunction has been implicated in both positive symptoms[90] and cognitive impairments[91]. Our observations of weakened cortical excitatory input to the striatum alongside preserved or elevated striatal excitatory coordination may be relevant to these symptom domains.

In contrast to decreased candidates, the increased glutamatergic candidates showed relatively low consensus in the biomedical literature, suggesting that these mechanisms remain unresolved. The low Alignment scores may partially reflect the framework’s current limitations in resolving indirect mechanistic relationships. For example, most refuting articles for the glutamatergic dysregulations discussed NMDA receptor hypofunction, and their nuances led the LLM to classify articles as contradictory to our observed glutamatergic changes. However, NMDA receptor hypofunction in GABAergic interneurons can produce downstream glutamatergic hyperactivity through disinhibition[92]. The pattern of refuting abstracts suggests that low literature consensus does not necessarily indicate a biological contradiction. Instead, the low consensus may reflect the multi-step mechanistic inference needed to relate these abstracts to our observations. In addition, some of the predictions, especially glycinergic interregional co-expression coordination changes and altered synaptic adhesion, have little prior literature evidence. Although our LLM-augmented literature screening provided a check of our candidates against existing literature, future optogenetic or conditional knockout experiments could test our hypotheses and elucidate specific interregional circuitry to confirm the directed relationships underlying the observed dysregulations.

Our analysis did not identify nominally significant dopaminergic dysregulation in patients with schizophrenia. Given the centrality of dopamine to schizophrenia pathophysiology and pharmacology[93], we consider that the absence of a dopaminergic signal in our results does not contradict current understanding. First, the dopaminergic abnormality in schizophrenia is predominantly presynaptic, affecting dopamine synthesis capacity and release rather than receptor or transporter expression[94, 95]. Since our method is built on the co-expression of genes, the method cannot precisely detect a functional abnormality in the synthesis and release of dopamine. Second, our sampled regions do not include the principal source regions of the dopaminergic system, the midbrain VTA and substantia nigra (SN). The VTA projects through mesolimbic and mesocortical dopaminergic pathways[96], and the SN projects through the nigrostriatal pathway[97]. Dopaminergic projections cannot be reconstructed because none of the integrated datasets include the VTA or SN, leaving only striatal and cortical dopaminergic coordination in the observable data. Third, the cohort included patients chronically treated with antipsychotics, untreated patients, and patients with unknown medication status. An analysis of the caudate nucleus transcriptome in a post-mortem cohort revealed substantial effects of antipsychotic treatment on gene expression in the region[41]. Chronic dopamine D2 receptor antagonism could account for the normalization or obscuring of diagnosis-related changes in dopaminergic co-expression coordination. More broadly, large-scale transcriptomic studies of schizophrenia have consistently identified synaptic, splicing, and cell-type signatures, rather than distinct dopaminergic transcriptional signals[98, 99]. Although dopamine is heavily emphasized in the pathophysiology of schizophrenia, the absence of a dopaminergic signal in our results is consistent with previous transcriptomic studies. Our results should not be interpreted as evidence against the dopaminergic hypothesis, because the hypothesis is based on functional measurements.

Along with neurotransmitter signaling abnormalities, we observed reduced coordination of the neurexin-neuroligin synaptic adhesion mechanism, which shapes neurotransmitter signaling in the synaptic cleft[100, 101]. These findings align with accumulating evidence for synaptic pathology in schizophrenia. NRXN1 deletions are established schizophrenia risk variants[102], reduced synaptic markers across cortical regions have been reported[103], and *in vivo* imaging has demonstrated reduced cortical synaptic densities[104, 105]. Our results add a directional and gene-specific dimension to cortical synaptic deficits by prioritizing distinct neurexin and neuroligin genes for each connection (NRXN3–NLGN2 for the cortical-to-striatal connection and NRXN2–NLGN3 for the striatal-to-cortical connection). Given that neurexin function is differentiated by alternative splicing and that individual isoforms specify distinct synapse types[106], this divergence suggests that the loss of synaptic adhesion is not uniform but region- and direction-specific. Both connections had low Relevancy. Although synaptic loss in schizophrenia is well established, its distribution across specific cortico-striatal pathways has received little direct attention.

The reduced cholinergic connection is also relevant to current therapeutic developments. Muscarinic receptor abnormalities have been implicated in schizophrenia for decades[107, 108]. These findings have motivated novel muscarinic therapies. Cobenfy combines xanomeline (an M1/M4-preferential muscarinic receptor agonist) and trospium chloride (a peripherally restricted muscarinic antagonist that alleviates the peripheral side effects of xanomeline). The drug was shown to be effective in a Phase 3 trial[109], and was subsequently approved by the FDA in 2024 as the first treatment for schizophrenia not acting through DRD2 blockade. However, the cholinergic evidence base is not uniform. The M4-selective positive allosteric modulator emraclidine failed in Phase 2 trials (Clinical trials NCT05227703 and NCT05227690), indicating that selective M4 engagement alone may not be sufficient and that therapeutic targeting of muscarinic receptors is still being investigated. Similarly, the partial α7 nicotinic receptor agonist DMXB-A has shown efficacy in cognitive enhancement in Phase 1 trials[110], but failed in Phase 2 trials[111, 112]. Our prioritized gene CHRNA7, which we identified as contributing to decreased cholinergic coordination, encodes the α7 nicotinic receptor. Our observation points to a mechanism that remains incompletely understood, and supports an emerging role for cholinergic signaling in schizophrenia.

Glycine enters our results in two ways. Two out of our 10 connections implicated glycinergic dysregulation, and a third – the intra-striatal glutamatergic CN → NAc connection – was driven partly by glycine-related receptor genes. Both glycinergic candidates returned a Relevancy score less than 0.0001. Unlike the synaptic adhesion candidates, which fall on direct or indirect cortico-striatal connections, glycinergic candidates have no established connection between the sampled regions. We therefore interpret them most cautiously. Glycine has two distinct roles in the central nervous system. It is a co-agonist, like D-serine, for the glutamatergic NMDA receptor. It is also the primary ligand for the inhibitory glycine receptor (GlyR), which is a chloride channel[18]. Our candidate dysregulations and prioritized genes implicate both roles. For the intra-striatal glutamatergic CN → NAc connection, we prioritized NMDA receptor subunit genes and associated glycine-related annotations. GRIN3A was prioritized exclusively for this connection. GRIN3A encodes NMDA receptor subunit 3A (GluN3A), which bears a glycine/serine modulatory site and assembles with subunit 1 (GluN1, encoded by the GRIN1 gene) to form a glycine-responsive receptor[113]. GluN3A alters the properties of NMDA receptors compared to classical receptors. When incorporated into the GluN1/GluN2 complex, GluN3A acts as a dominant-negative subunit that makes the receptor less permeable to Ca^2+^ and relatively insensitive to Mg^2+^-induced block[114]. Thus, connection-specific prioritization of GRIN3A helps explain downstream enrichment of glycine-related biological terms, reflecting the neurochemical properties of GluN3A-containing receptors. These properties shape how the glutamatergic coordination should be interpreted, as the increased coordination does not necessarily indicate enhanced NMDA receptor function, particularly when a subunit with these properties is involved. Together with the reduction in glycinergic coordination between other regions, the pattern is consistent with attenuated NMDA receptor signaling accompanied by excessive excitatory coordination, and is compatible with the NMDA receptor hypofunction hypothesis.

The prioritization of glycine receptors supports another role of glycine, as an inhibitory neurotransmitter acting through chloride conductance. Two glycinergic candidate dysregulations were driven by the inhibitory GlyR subunit GLRB. GlyR-mediated inhibitory signaling depends on the transmembrane chloride gradient, established by cation-chloride cotransporters (CCCs), mainly KCC2 (SLC12A5) and NKCC1 (SLC12A2)[115]. This dependency is relevant to our prioritization of the contributing gene NLGN2 in synaptic adhesion deficits. NLGN2 is associated with the dysfunction of KCC2[116], and the gene is also essential for GABAergic neurotransmission and synaptic development[117]. The impairment of synaptic adhesion along with an abnormal transmembrane chloride gradient could contribute to dysregulated GlyR and GABA receptor signaling[118]. Although the two roles of glycine are reflected in our results, we also note that CCCs were not included in our curated gene list. Therefore, the mechanisms proposed here are hypotheses that are consistent with the direction of our observations and supported by the literature.

Functional inhibitory glycine receptors and transport mechanisms do exist locally in striatal regions and the hippocampus. Glycine receptor currents have been reported in striatal medium spiny neurons and in broader forebrain regions[119, 120], and morphologically identified glycinergic synapses have been described within the hippocampus[121]. However, there are no established intra-striatal or striato-hippocampal monosynaptic glycinergic projections. Glycine signaling has been proposed as a therapeutic target beyond dopaminergic receptor antagonism, by inhibiting glycine transporter 1 (GlyT1) to potentiate NMDA receptor co-agonism[18]. Three GlyT1-inhibitory compounds have been tested clinically: bitopertin, iclepertin, and sarcosine (N-methylglycine)[122–124]. However, this rationale has not translated into clinical benefit. Bitopertin and iclepertin both produced encouraging Phase 2 trial results but failed in Phase 3 trials[125, 126], and sarcosine add-on treatment trials reported mixed effects[124]. These trials tested systemic GlyT1 inhibition, whereas our findings include changes in glycinergic co-expression coordination within specific connections. Our results do not allow us to determine whether regional specificity of this kind contributes to the failure of global glycine transporter inhibition. We therefore present our findings as evidence of an underexplored circuit-level relationship, but not as support for a therapeutic strategy that recent trial evidence has weakened.

Our method rests on the assumption that correlated expression of presynaptic genes in one region and postsynaptic genes in another region reflects coordinated signaling capacity between the two regions. This premise is supported by evidence that interregional gene co-expression covaries with anatomical and functional connectivity in the human and rodent brain[26, 127]. Our macaque benchmarking result also partially supports this assumption. However, interregional co-expression across individuals is a statistical association, not a physiological measurement. That association may reflect shared interindividual variation, which likely arises from coordinated signaling or from shared systemic influences, developmental trajectories, medication status, or covarying cellular composition across regions[128]. Furthermore, there has been debate about whether the association between gene co-expression and functional brain networks is an artifact of uncontrolled proximity between brain regions[129], or whether it reflects genuine coordination[130]. Our design of abundance-weighting and downstream directional consistency filtering requires that inferred changes not contradict source-region synthesis capacity. Our label permutation analysis calibrates the difference of each connection against its own null distribution. However, neither addresses the underlying inferential limitation. Therefore, we describe our findings as changes in co-expression coordination, and reserve circuit-level language for interpretation.

We also acknowledge that our analysis was restricted to specific brain regions available through the dataset, potentially missing key nodes in the broader schizophrenia network. Consequently, there are several limitations in interpreting our candidate dysregulations. Our data do not measure thalamic participation, since that region was not sampled. The CSTC framing is therefore an interpretive scaffold for our observations. For example, GABAergic NAc output only reaches the cortex indirectly, through the ventral pallidum and thalamic nuclei[131]. Although descending cortical output to the striatum has been well established, striatal outputs to cortex are indirect polysynaptic pathways mediated through relay regions. Our findings should be interpreted as aggregated indirect coordination changes within the CSTC loop, not as evidence of direct synaptic pathways. The same caution also applies to interpreting cholinergic and glycinergic dysregulations. Although our study suggested a candidate striato-hippocampal cholinergic connection, no direct monosynaptic connection between these regions has been reported. Cholinergic interneurons are present in both the striatum and the hippocampus, but cholinergic projection neurons are mainly located in the brainstem or basal forebrain[132]. Hippocampal cholinergic synthesis genes fell below our expression threshold, resulting in an estimated acetylcholine abundance of zero. Glycine receptors are primarily found within the brainstem and spinal cord[133], and no long-range interregional glycinergic projections between the sampled regions have been established. Direct glutamatergic connections between CN and NAc are not established, despite their adjacency. Connections between the regions are relayed through midbrain regions[134]. For these candidates, simultaneous attenuation of several neurotransmitter-specific connections on the same directed edge might reflect shared co-regulatory influence on co-expression coordination between presynaptic and postsynaptic regions. The co-regulation likely results from multi-regional circuit-level relationships, or common regulatory influences on both regions originating from other factors that we could not measure. The directionality of our connections is derived from the presynaptic and postsynaptic classification of participating genes, and transcriptional correlation alone cannot distinguish genuine coordination from these alternatives. Our reconstruction of the CSTC circuit is therefore partial, and including additional relevant regions in future work would help us understand complex dysregulations across a wider set of regions.

Several intrinsic limitations arise from using bulk transcriptomic data, which are aggregated expression profiles across all cell types within a tissue sample. Transcriptomic data are static, and gene expression does not always correlate linearly with protein levels or synaptic efficacy. Without using single-cell-resolution transcriptomic data, the contributions of individual neuron subtypes and glial cells cannot be resolved. Although post-mortem tissue is the primary resource for multi-regional human transcriptomic profiling, it could introduce confounding variables. For example, chronic antipsychotic treatment results in altered levels of several signaling-related proteins[41, 135]. Although our analysis included background correction, a large sample size, and a directional consistency filter intended to mitigate false-positive findings, some observed dysregulations may partially reflect compensatory responses to treatment. Further studies using other modalities, such as longitudinal sampling, scRNA-seq, and single-nucleus RNA-seq, would help reconstruct temporal interregional changes in connectivity with respect to distinct cell types. Stratifying medication status in the human cohort would help isolate the primary pathological mechanism of schizophrenia from treatment-influenced compensatory artifacts.

## Conclusions

Using transcriptomic data from five brain regions, we constructed an “interregional molecular connectome” estimated from coordination between presynaptic and postsynaptic neurotransmission-related genes. Our estimation pipeline predicted macaque anatomical connectivity better than whole-transcriptome and random gene co-expression. We identified a coherent pattern of altered interregional co-expression coordination that is consistent with the CSTC circuit, including bidirectional dysregulation in cortico-striatal circuitry involving multiple neurotransmitter systems. The changes include reduced synaptic adhesion coordination and glycinergic candidate dysregulations, each of which could destabilize functional neural circuits within the affected regions. These results are exploratory, and we treat our observations as hypothesis-generating candidates and interpret them against established anatomical relationships. By integrating the computational reconstruction of the neurotransmission connectivity network with calibrated LLM-augmented literature screening, we annotated our findings with quantified literature support. We provide an “interregional molecular connectome” that supports a unified multi-regional, multi-neurotransmitter perspective of schizophrenia as a systemic failure of neurotransmission, and points to molecular mechanisms. The relatively underexplored glycinergic and synaptic adhesion candidates could provide direct targets for experimental follow-up. Further investigation of these candidates could help link genetic risk, molecular expression, endophenotypes, and clinical phenotypes. We also expect that this study will offer an approach that can be applied to other psychiatric disorders characterized by multi-regional dysregulations.

## List of abbreviations

GABA: γ-aminobutyric acid
AMPA: α-amino-3-hydroxy-5-methyl-4-isoxazolepropionic acid
NMDA: N-methyl-D-aspartate
CCC: Cation-chloride cotransporter
DRD2: Dopamine receptor D2
GlyR: Glycine receptor
PFC: Prefrontal cortex
CC: Cingulate cortex
HP: Hippocampus
CN: Caudate nucleus
NAc: Nucleus accumbens
VISp: Primary visual cortex
ALM: Anterior lateral motor cortex
VTA: Ventral tegmental area
SN: Substantia nigra
CSTC: Cortico-striato-thalamo-cortical circuit
SMRI: Stanley Medical Research Institute
MA: Microarray
RNA-seq: RNA sequencing
scRNA-seq: Single-cell RNA sequencing
CPM: Counts per million
SVA: Surrogate variable analysis
PCC: Pearson correlation coefficient
CDF: Cumulative distribution function
GO: Gene Ontology
ROC: Receiver operating characteristic curve
PR: Precision-recall curve
auROC: Area under the ROC curve
auPRC: Area under the PR curve
CI: Confidence interval
BH: Benjamini-Hochberg procedure
ρ: Spearman’s rank correlation
CV: Coefficient of variation
LLM: Large language model
RAG: Retrieval-augmented generation
NLI: Natural language inference
GPU: Graphics processing unit

## Declarations

### Ethics approval and consent to participate

Not applicable. This study analyzed publicly available datasets (GEO series GSE53987, GSE80655, GSE180092, GSE115746, and GSE185862) and restricted-access datasets (dbGaP phs000979.v3.p2, phs003495.v1.p1.c1) together with data from the Stanley Medical Research Institute (SMRI). All datasets were obtained with approval from the relevant ethics committees for the original projects. No new human participants or identifiable human data were used.

### Consent for publication

Not applicable. No individual-level participant information, images, or identifiable data are reported.

## Availability of data and materials

### Data availability

Transcriptomic data: GSE53987, GSE80655, GSE180092, GSE115746, and GSE185862 are publicly available through the GEO database; BrainSeq PFC and HP data are publicly available through “BrainSeq Phase 2” website (https://eqtl.brainseq.org/phase2/); HBCC data and BrainSeq CN data are available upon request under dbGaP (HBCC Accession: phs000979.v3.p2 and BrainSeq CN Accession: phs003495.v1.p1.c1); Three SMRI datasets (Array Collection, New Collection, Neuropathology Consortium) are available upon request to the SMRI research advisor. Macaque tract-tracing data ‘Organized CoCoMac’ and ‘TheVirtualBrain’ are available through their original publications. Mouse retrograde tracer data are available through publication by Tasic *et al*. Other resources, including gene set data, are available through their original works.

### Code availability

All code used in this work is available through a GitHub repository (https://github.com/bisl-hub/MolConnectome) under the GPL-3.0 license.

## Competing interests

The authors declare that they have no competing interests.

## Funding

This research was supported by the Bio&Medical Technology Development Program of the National Research Foundation (NRF) funded by the Korean government (MSIT) (No. RS-2025-16063391).

## Authors’ contributions

OK: Conceptualization, Methodology, Investigation, Software, Visualization, Writing – Original Draft

UK: Methodology, Software, Writing – Original Draft

MJW: Writing – Review & Editing, Resources, Data Curation

SK: Supervision, Writing – Review & Editing, Resources, Data Curation

DL: Supervision, Writing – Review & Editing, Funding acquisition

All authors reviewed and approved the final manuscript.

## Acknowledgements

We thank the Stanley Medical Research Institute for providing access to post-mortem brain tissue data.

## Supplements

### Supplementary Figures

**Supplementary Fig. S1.**
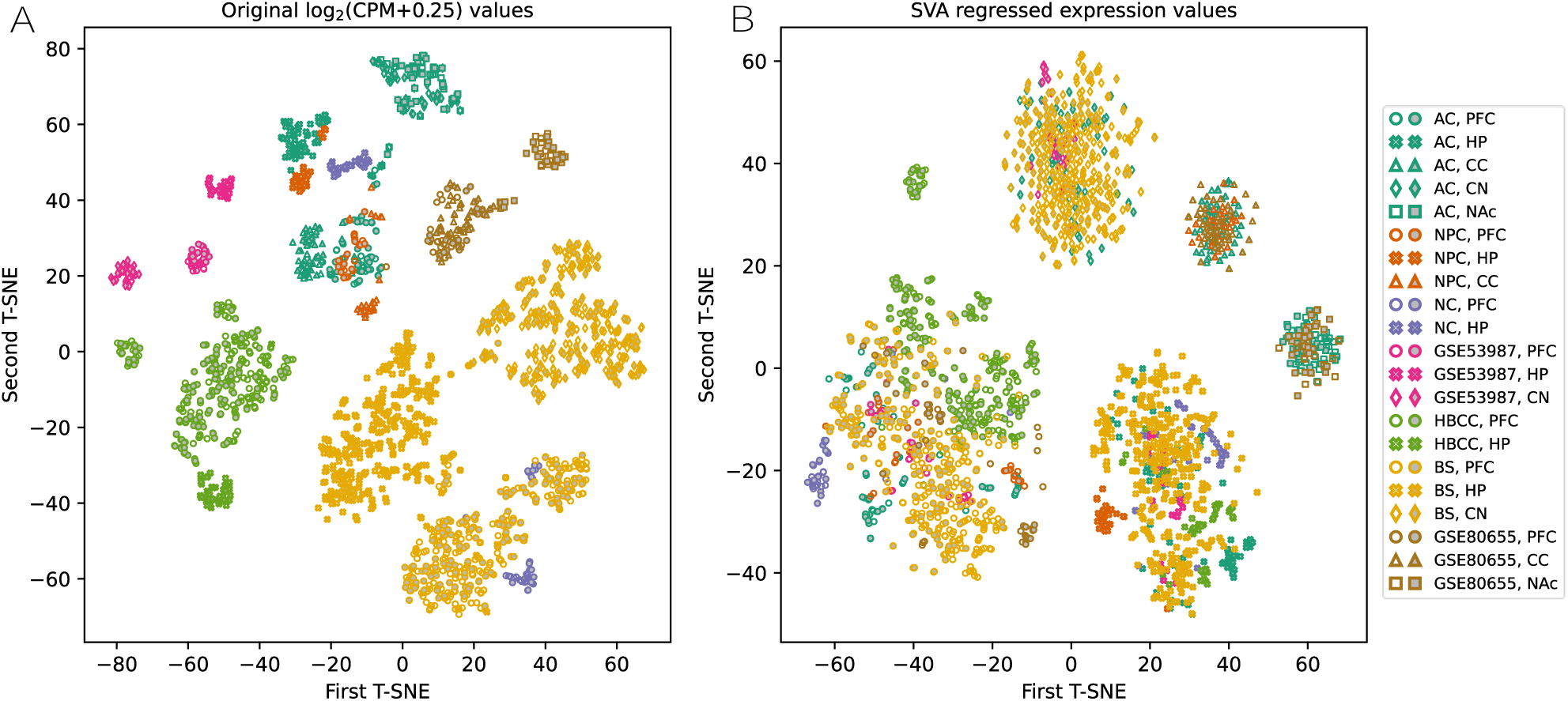
Human expression profiles before and after surrogate variable analysis. t-SNE dimensionality embeddings of the 1,884 human samples (**A**) before and (**B**) after SVA corrections. Color denotes dataset and shape denotes brain region. Samples from patients with schizophrenia are filled grey and samples from controls are hollow. Dataset abbreviations: AC: Array Collection; NPC: Neuropathology Consortium; NC: New Collection; HBCC: Human Brain Collection Core; BS: BrainSeq. Region abbreviations: PFC: prefrontal cortex; CC: cingulate cortex; HP: hippocampus; CN: caudate nucleus; NAc: Nucleus accumbens.

**Supplementary Fig. S2.**
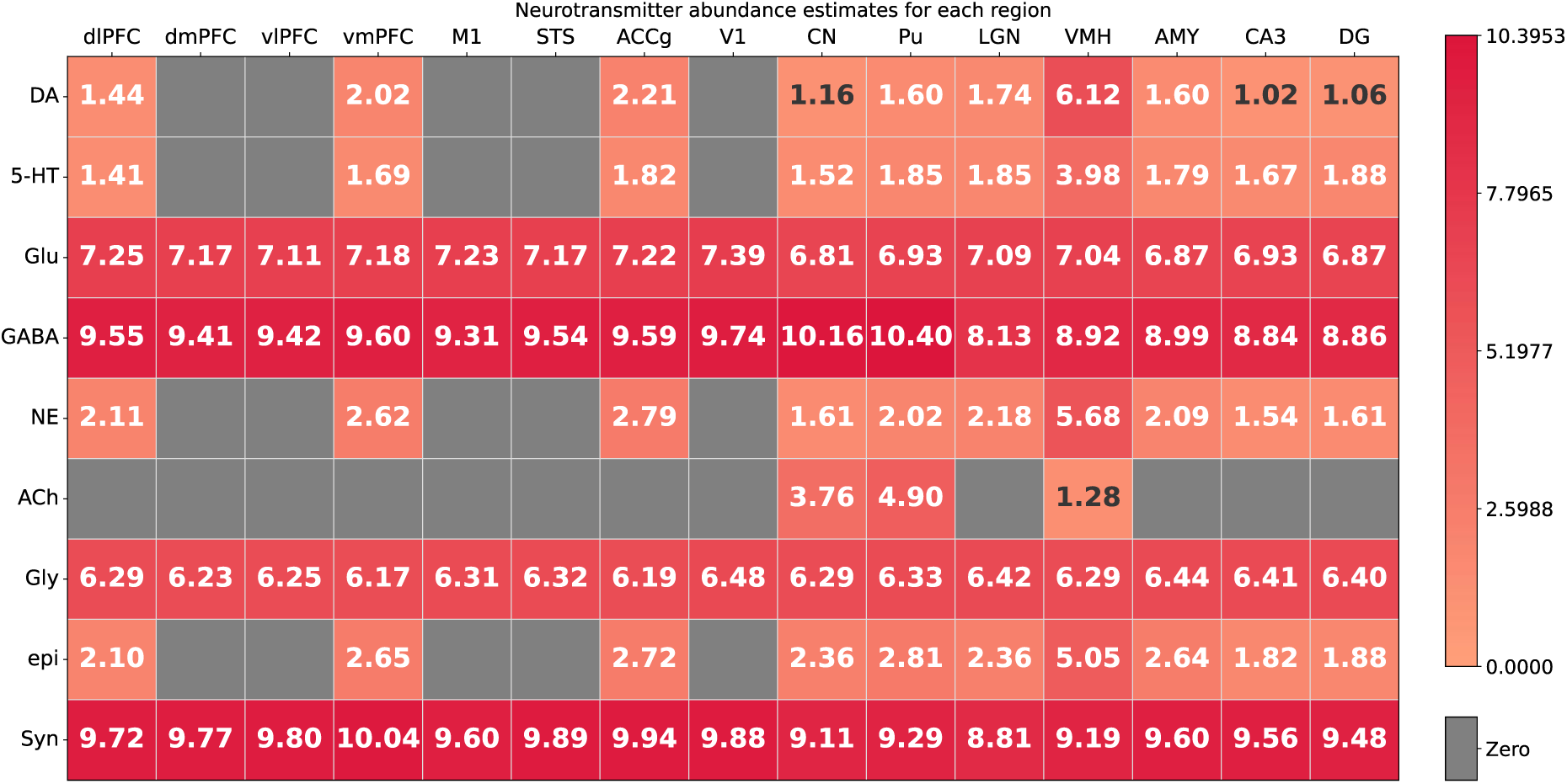
Estimated neurotransmitter abundance in 15 macaque brain regions. Abundance of each neurotransmitter was estimated for each sample from the expression of its core synthetic enzymes with “AND”-’OR” logic adapted from “NeuronChat”. Synaptic adhesion abundance is the mean expression of neurexin genes (NRXN1, NRXN2, NRXN3). Values are means across the 36 animals, which were used to weight connectivity strengths in the macaque benchmark. A zero estimate means that one or more core synthetic enzymes fell below the expression threshold. Region abbreviations: dlPFC: dorsolateral prefrontal cortex; dmPFC: dorsomedial prefrontal cortex; vlPFC: ventrolateral prefrontal cortex; vmPFC: ventromedial prefrontal cortex; M1: primary motor cortex; STS: superior temporal sulcus; ACCg: anterior cingulate cortex; V1: primary visual cortex; CN: caudate nucleus; Pu: putamen; LGN: lateral geniculate nucleus; VMH: ventromedial hypothalamus; AMY: amygdala; CA3: cornu ammonis 3; DG: dentate gyrus. Neurotransmitter abbreviations: DA: dopamine; 5-HT: serotonin; Glu: glutamate; GABA: γ-aminobutyric acid; NE: norepinephrine; ACh: Acetylcholine; Gly: glycine; epi: epinephrine; Syn: synaptic adhesion.

**Supplementary Fig. S3.**
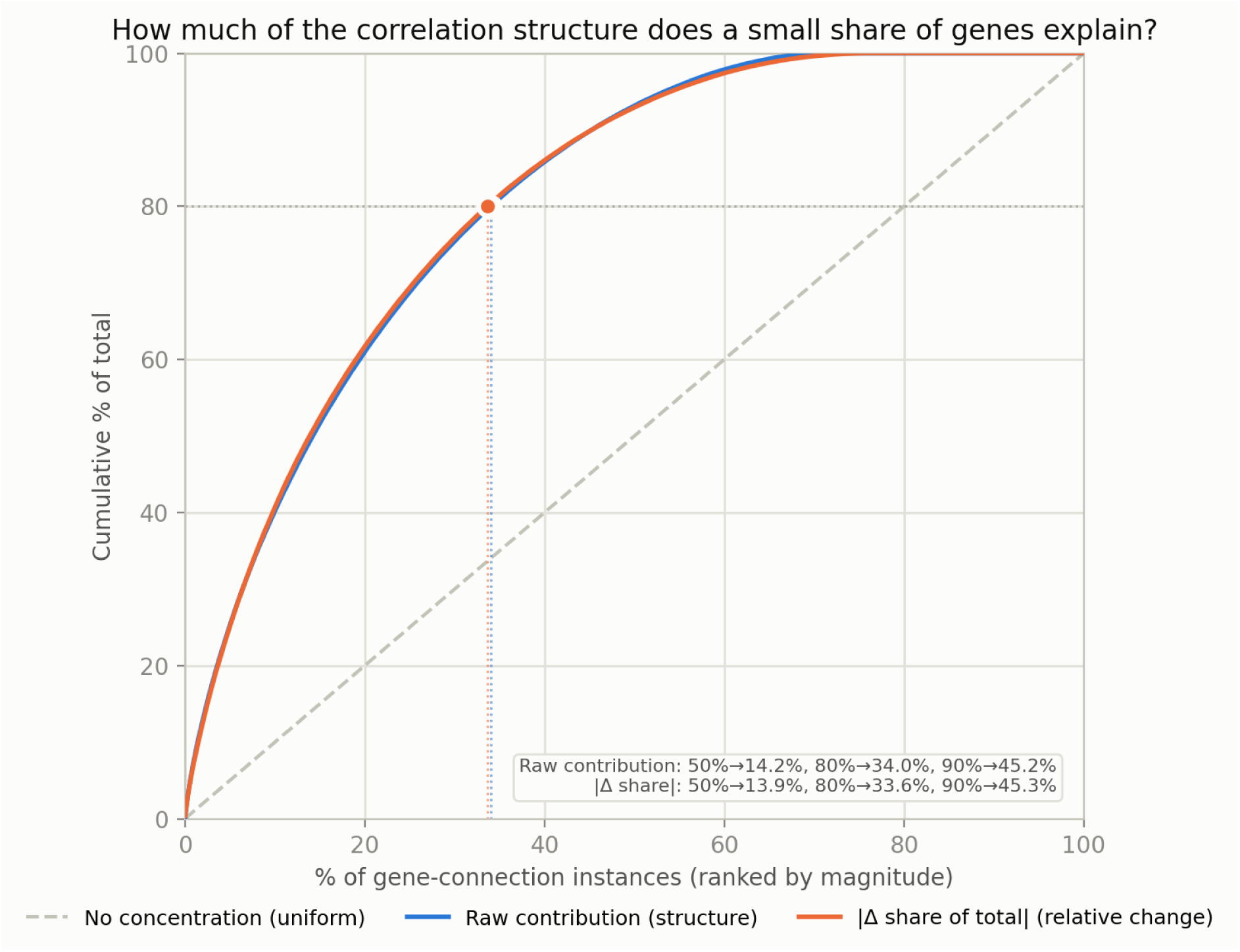
Cumulative share of correlation contributions across ranked genes. The correlation contribution of each gene was calculated as the sum of its PCC over all gene pairs in the connection. Genes were ranked by contribution. The curve shows the cumulative fraction of the total contribution against the top-percentile cutoff. The top 30% of genes, selected as prioritized genes, account for nearly 80% of the total correlation contributions.

**Supplementary Fig. S4.**
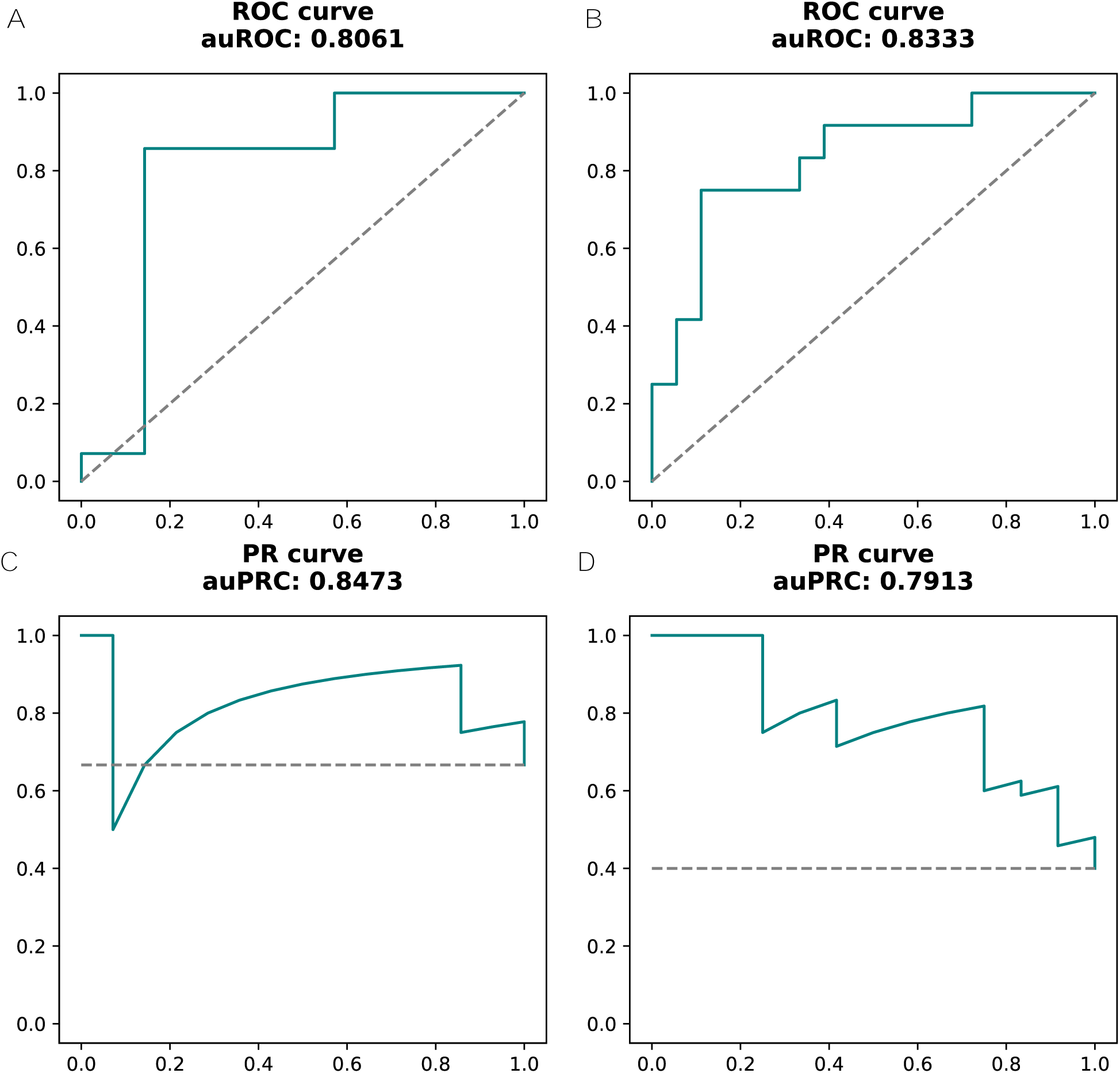
Performance of the “NeuronChat” method on pseudo-bulk mouse transcriptomes. Mouse scRNA-seq data were aggregated into pseudo-bulk profiles per sample, processed with our preprocessing pipeline, and analysed with the original “NeuronChat” package. Predicted connection strengths were benchmarked against retrograde injection data. (**A**) ROC curve for the VISp model. (**B**) ROC curve for the ALM model. (**C**) PR curve for the VISp model. (**D**) PR curve for the ALM model. The original study reported auROC = 0.8316 and auPRC = 0.9153 for the VISp model; auROC = 0.7639 and auPRC = 0.7677 for the ALM model. Abbreviations: ALM: anterior lateral motor cortex; VISp: primary visual cortex; ROC curve: Receiver Operating Characteristic curve; PR curve: Precision-Recall curve; auROC: area under the ROC curve; auPRC: area under the PR curve.

**Supplementary Fig. S5.**
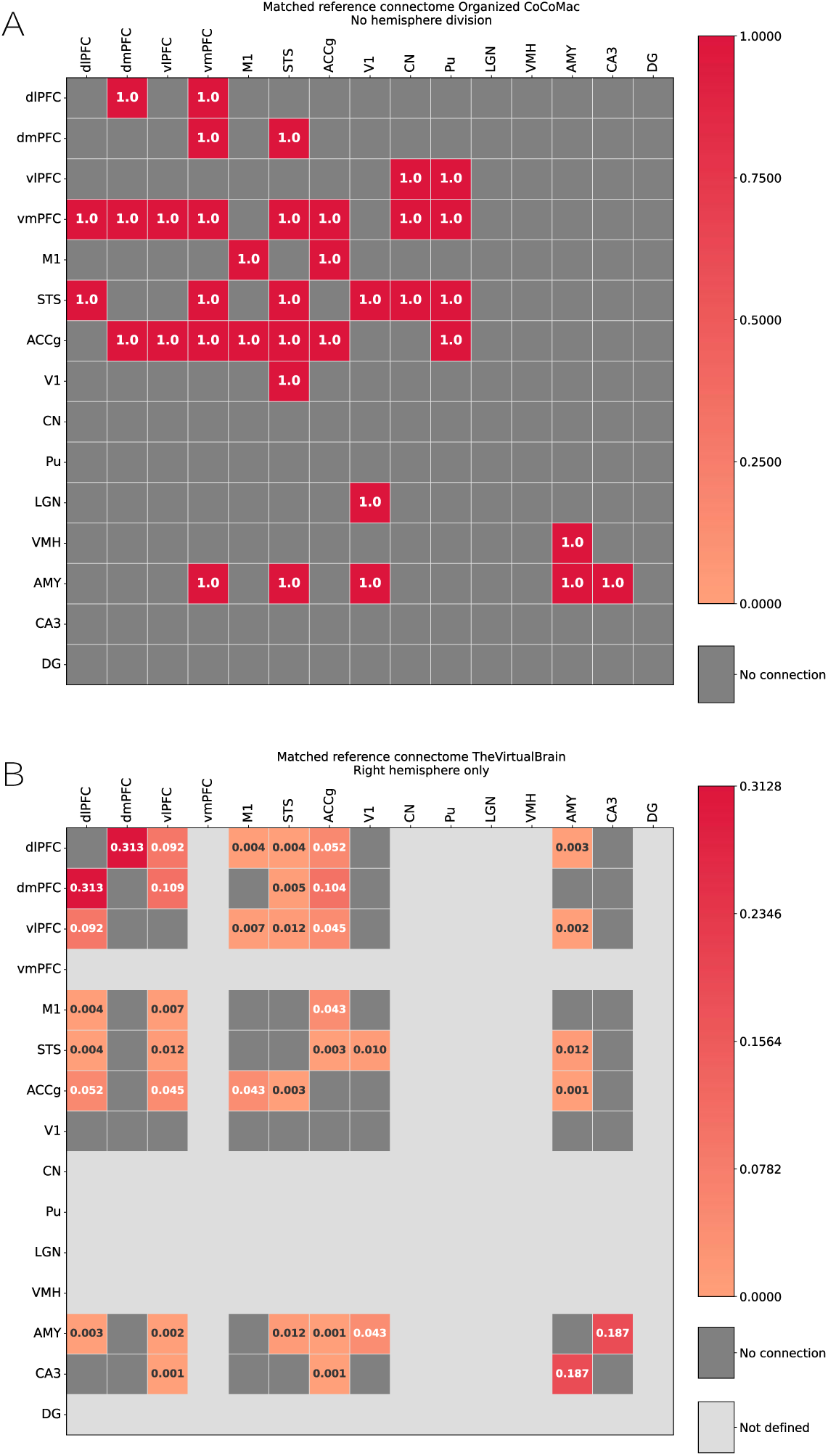
Macaque reference connectomes derived from the CoCoMac tract-tracing database. Directed connections among the 15 transcriptomic regions of GSE180092 after mapping onto each connectome’s parcellation. Rows are source regions, and columns are target regions. (**A**) “Organized CoCoMac” connectome. (**B**) **“**TheVirtualBrain” connectome. Region abbreviations: dlPFC: dorsolateral prefrontal cortex; dmPFC: dorsomedial prefrontal cortex; vlPFC: ventrolateral prefrontal cortex; vmPFC: ventromedial prefrontal cortex; M1: primary motor cortex; STS: superior temporal sulcus; ACCg: anterior cingulate cortex; V1: primary visual cortex; CN: caudate nucleus; Pu: putamen; LGN: lateral geniculate nucleus; VMH: ventromedial hypothalamus; AMY: amygdala; CA3: cornu ammonis 3; DG: dentate gyrus.

**Supplementary Fig. S6.**
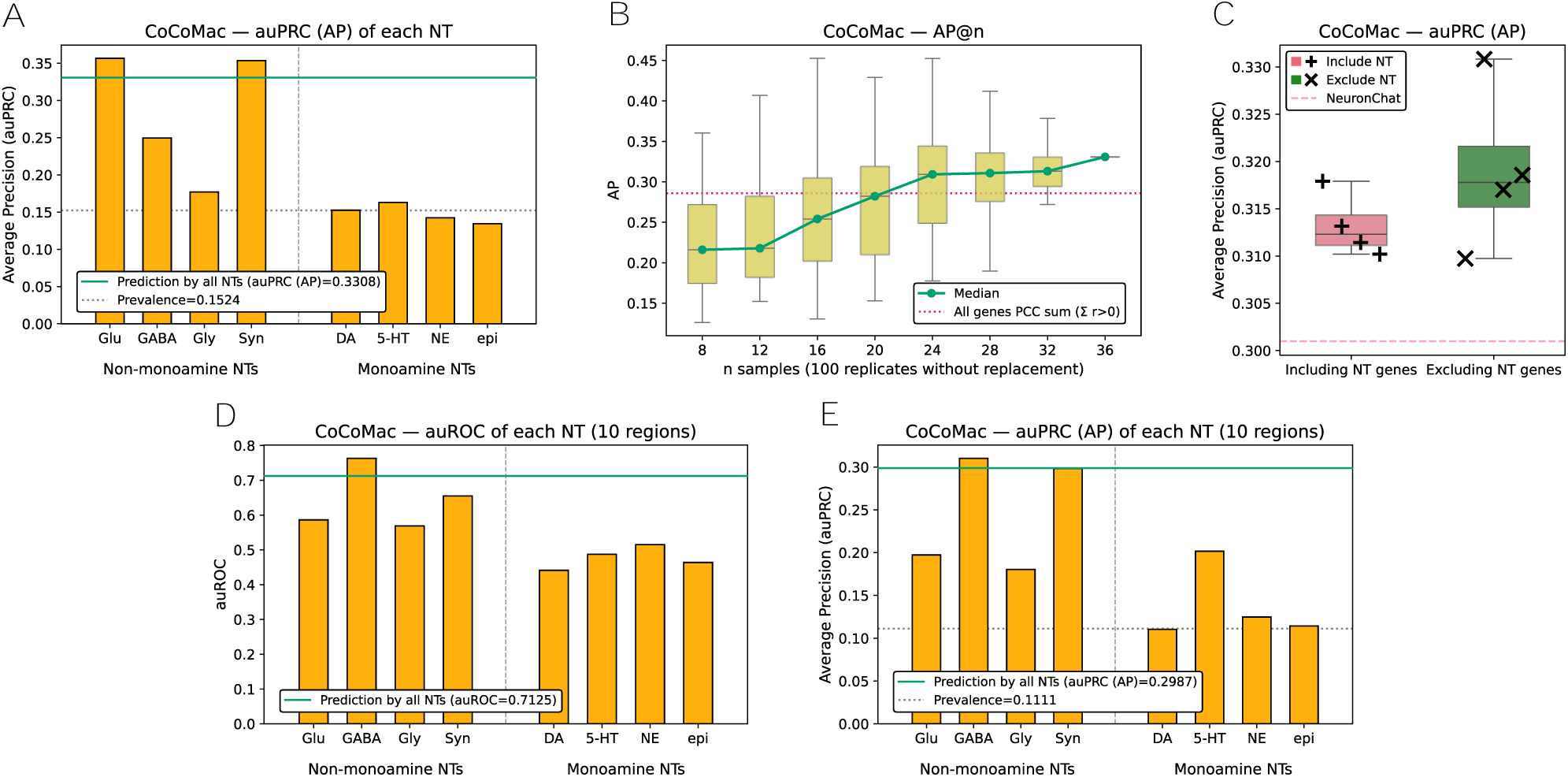
Additional benchmarking results against the Organized CoCoMac reference connectome. Panels (A)-(C) complement Figs. 2E-G. (**A**) auPRC for each seven neurotransmitter systems and synaptic adhesion across 15 regions. Acetylcholine was excluded. (**B**) auPRC across subsample sizes (n = 8 to 36 animals in steps of 4; 100 subsamples drawn without replacement per size), as in Fig. 2F. (**C**) auPRC across the eight background gene panel sets shown in Fig. 2G. (**D**) auROC and (**E**) auPRC for each of seven neurotransmitter systems and synaptic adhesion, with the predicted connections and the reference connectome both restricted to the 10 regions with non-zero estimated abundance.

**Supplementary Fig. S7.**
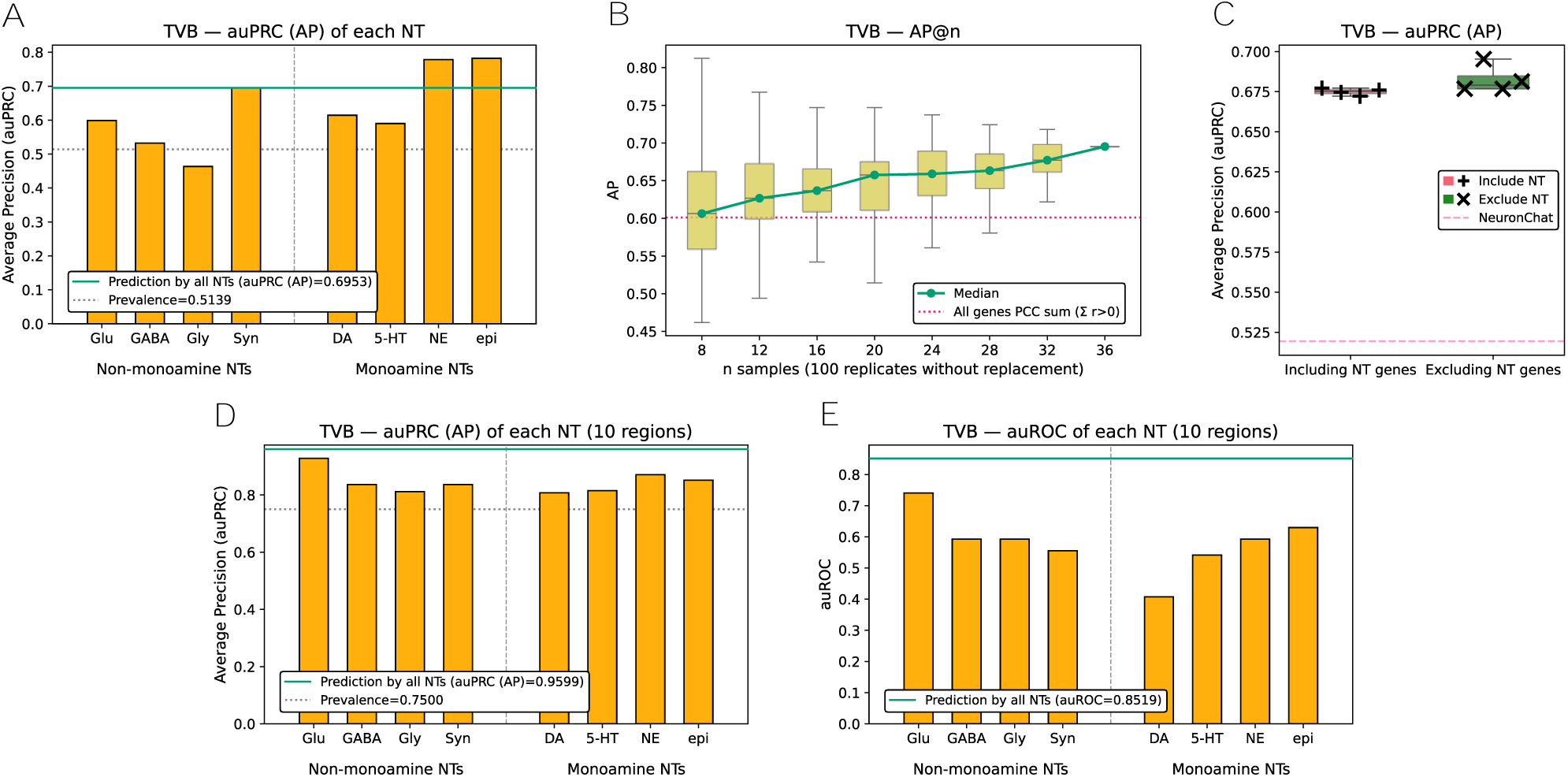
Additional benchmarking results against TheVirtualBrain reference connectome. Panels (A)-(C) complement Figs. 3E-G. (**A**) auPRC for each seven neurotransmitter systems and synaptic adhesion across 15 regions. Acetylcholine was excluded. (**B**) auPRC across subsample sizes (n = 8 to 36 animals in steps of 4; 100 subsamples drawn without replacement per size), as in Fig. 3F. (**C**) auPRC across the eight background gene panel sets shown in Fig. 3G. (**D**) auROC and (**E**) auPRC for each of seven neurotransmitter systems and synaptic adhesion, with the predicted connections and the right-hemisphere reference connectome both restricted to the 10 regions with non-zero estimated abundance.

## Supplementary Information

**Supplementary information S1** Detailed information about gene list curation.

We curated genes using the information from these four previous works, resulting in a total of 585 genes that are associated with neurotransmission, which are classified by their functionalities and associated neurotransmitter systems. The detailed list of curated genes, their membership, and category classification is in Supplementary Table S2.

### 1) The Allen Human Brain Atlas (AHBA) gene list

The AHBA manually curated genes that are related to neurotransmission and listed them. The original purpose of this gene list was to use the listed genes in the in situ hybridization analysis. The original gene list is a table with the following columns: 1) gene symbol, 2) Entrez ID for the gene, 3) description of the gene (i.e., full name), 4) regional enrichment, 5) associated neurotransmitter system, 6) gene family (i.e., their functionalities), and 7) marker type. In this table, we used the gene symbol, its associated neurotransmitter system, and the gene family columns. The original gene classification is available through the AHBA official website (https://community.brain-map.org/t/documentation-human-brain-atlas/2879 in the ‘Gene list’ file).

The neurotransmitter system column originally consisted of 11 different neurotransmitters: acetylcholine, adenosine, cannabinoid, dopamine, epinephrine/norepinephrine, GABA, glutamate, glycine, histamine, neuropeptide, and serotonin. In our study, we only focused on small molecular neurotransmitters and neurotransmitter systems with a sufficient number of related genes, which thus excludes the neurotransmitter systems of 1) adenosine, 2) cannabinoid, 3) histamine, and 4) neuropeptide. This resulted in a total of eight different neurotransmitter systems to focus on. The genes that are related to the four excluded neurotransmitter systems or not related to any of the 11 neurotransmitters (that is, the field was empty) were considered as the “other” system. These genes were included in the 585 superset, but not considered further in our manual curation.

The “gene family” column originally consisted of 13 different values: cell adhesion, cytoskeletal protein, GPCR, ion channel, metabolic enzyme, other intracellular, other membrane protein, peptide ligand, secreted protein, signal transduction, synaptic protein, transcription factor, and transporter. We then chose to assign the genes to Category I if the gene is associated with any different steps of the signal transduction process and to Category II if the gene is not directly related to the signal transduction process. This resulted in the genes classified under the gene families 1) GPCR, 2) ion channel, and 3) signal transduction were assigned to Category I, and the rest to Category II.

### 2) Original work by Bo et al. (2023)

Bo et al. analyzed the expression levels of genes that are associated with neurotransmission from multiple regions of the macaque brain. Similar to the AHBA gene list, they comprehensively curated the genes with their associated neurotransmitter systems and the functionalities of the genes. The neurotransmitter systems were the same as the eight neurotransmitters selected in the AHBA gene list. In their work, they classified the functionalities of the gene into five different classes: receptor, breakdown, synthesis, reuptake, and packaging. We assigned the genes that are marked as receptors to Category I and the remaining genes to Category II, as with the AHBA gene list.

### 3) The Reactome database

The Reactome database is a publicly available, manually curated pathway database. Many biological functions are organized in a tree-like shape, providing the genes and molecules that are related to the functions and sub-categories. We first selected the largest pathway term “Neuronal System” (Reactome ID: R-HSA-112316) in our gene list curation. The “Neuronal System” pathway consists of several different branch pathways, and the branches recursively consist of sub-branches with more specific functionalities. We recursively inspected all the possible sub-branches of the “Neuronal system” pathway and manually decided which pathways could be assigned to any of the nine neurotransmitter systems and two categories in each neurotransmitter system. We further searched for the names of each neurotransmitter and manually listed the Reactome terms related to the metabolism and signal transduction of each neurotransmitter that are not included in the “Neuronal system” term (which were mainly from the top-level term “Metabolism” and “Signal Transduction”). A detailed list of this curation is in the following table. We used the V94 version of the Reactome database (accessed 2025-09-14), and their whole pathway information with entities included is available through their website (https://reactome.org/download-data).

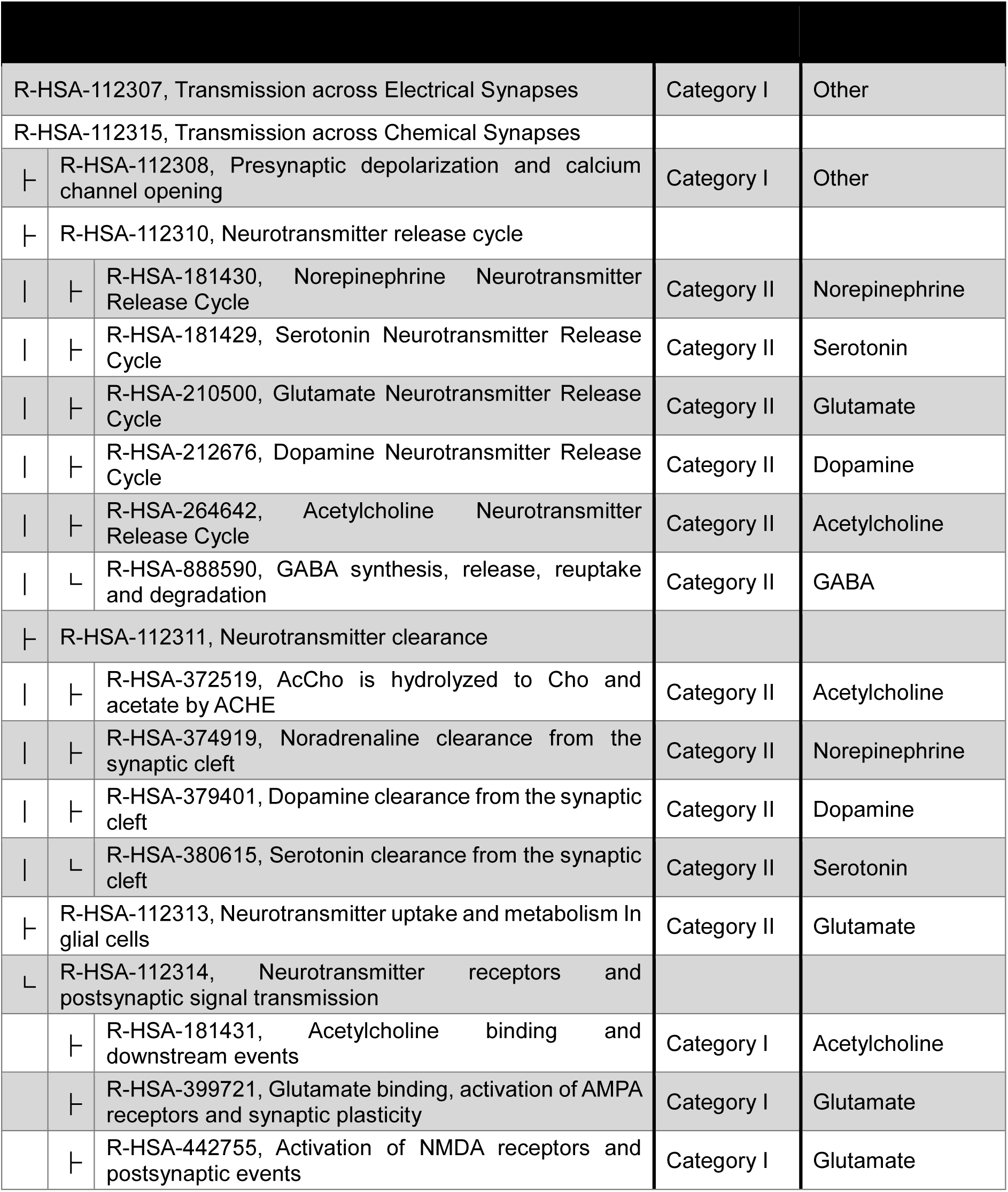
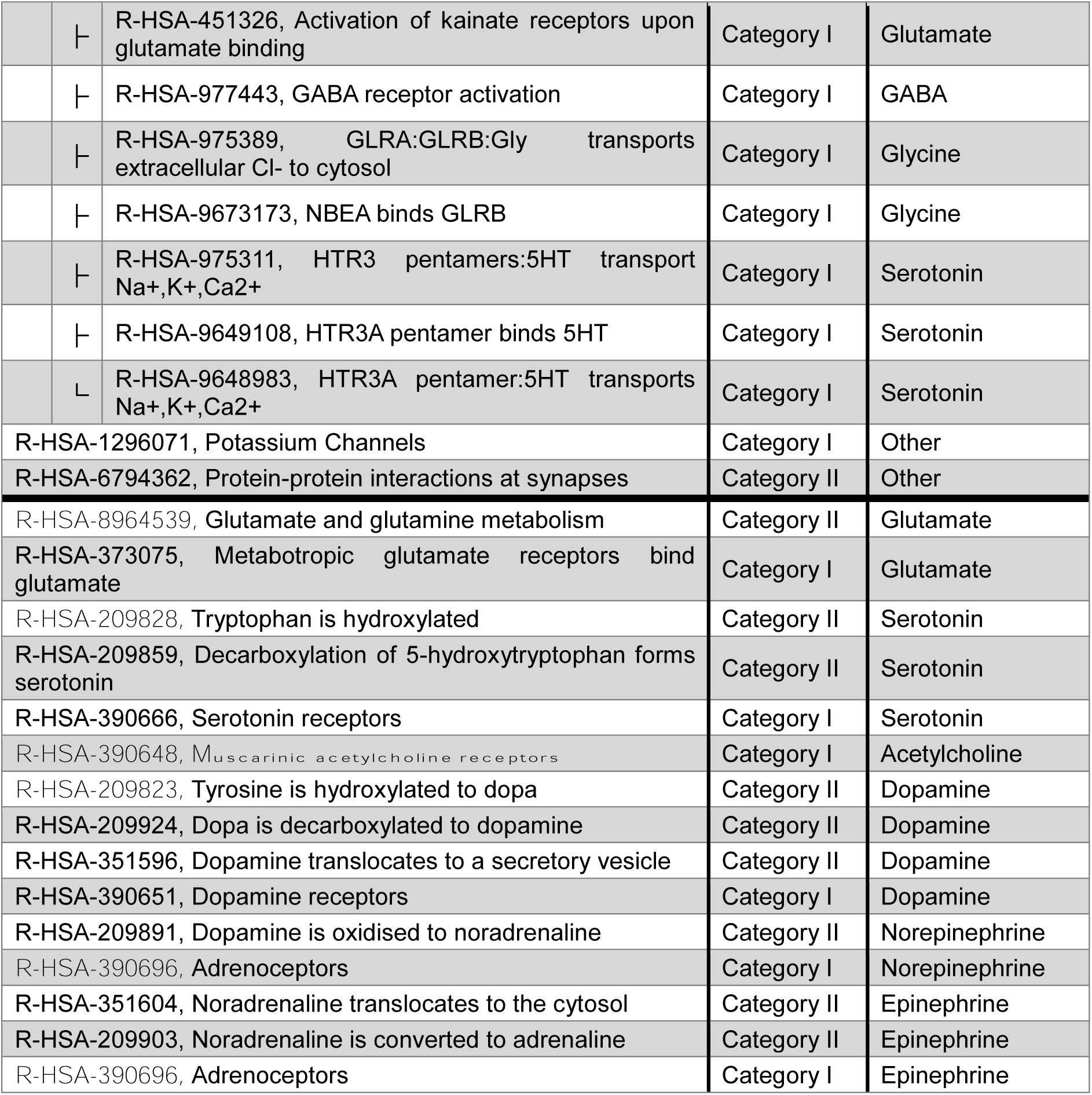

| R-HSA-112307, Transmission across Electrical Synapses |  |  | Category I | Other |
| --- | --- | --- | --- | --- |
| R-HSA-112315, Transmission across Chemical Synapses |  |  |  |  |
| └ | R-HSA-112308, Presynaptic depolarization and calcium channel opening |  | Category I | Other |
| └ | R-HSA-112310, Neurotransmitter release cycle |  |  |  |
|  | └ | R-HSA-181430, Norepinephrine Neurotransmitter Release Cycle | Category II | Norepinephrine |
|  | └ | R-HSA-181429, Serotonin Neurotransmitter Release Cycle | Category II | Serotonin |
|  | └ | R-HSA-210500, Glutamate Neurotransmitter Release Cycle | Category II | Glutamate |
|  | └ | R-HSA-212676, Dopamine Neurotransmitter Release Cycle | Category II | Dopamine |
|  | └ | R-HSA-264642, Acetylcholine Neurotransmitter Release Cycle | Category II | Acetylcholine |
|  | └ | R-HSA-888590, GABA synthesis, release, reuptake and degradation | Category II | GABA |
| └ | R-HSA-112311, Neurotransmitter clearance |  |  |  |
|  | └ | R-HSA-372519, AcCho is hydrolyzed to Cho and acetate by ACHE | Category II | Acetylcholine |
|  | └ | R-HSA-374919, Noradrenaline clearance from the synaptic cleft | Category II | Norepinephrine |
|  | └ | R-HSA-379401, Dopamine clearance from the synaptic cleft | Category II | Dopamine |
|  | └ | R-HSA-380615, Serotonin clearance from the synaptic cleft | Category II | Serotonin |
| └ | R-HSA-112313, Neurotransmitter uptake and metabolism In glial cells |  | Category II | Glutamate |
| └ | R-HSA-112314, Neurotransmitter receptors and postsynaptic signal transmission |  |  |  |
|  | └ | R-HSA-181431, Acetylcholine binding and downstream events | Category I | Acetylcholine |
|  | └ | R-HSA-399721, Glutamate binding, activation of AMPA receptors and synaptic plasticity | Category I | Glutamate |
|  | └ | R-HSA-442755, Activation of NMDA receptors and postsynaptic events | Category I | Glutamate |

|  |  |  |  |
| --- | --- | --- | --- |
| ┆ | R-HSA-451326, Activation of kainate receptors upon glutamate binding | Category I | Glutamate |
| ┆ | R-HSA-977443, GABA receptor activation | Category I | GABA |
| ┆ | R-HSA-975389, GLRA:GLRB:Gly transports extracellular Cl- to cytosol | Category I | Glycine |
| ┆ | R-HSA-9673173, NBEA binds GLRB | Category I | Glycine |
| ┆ | R-HSA-975311, HTR3 pentamers:5HT transport Na+,K+,Ca2+ | Category I | Serotonin |
| ┆ | R-HSA-9649108, HTR3A pentamer binds 5HT | Category I | Serotonin |
| ┆ | R-HSA-9648983, HTR3A pentamer:5HT transports Na+,K+,Ca2+ | Category I | Serotonin |
|  | R-HSA-1296071, Potassium Channels | Category I | Other |
|  | R-HSA-6794362, Protein-protein interactions at synapses | Category II | Other |
|  | R-HSA-8964539, Glutamate and glutamine metabolism | Category II | Glutamate |
|  | R-HSA-373075, Metabotropic glutamate receptors bind glutamate | Category I | Glutamate |
|  | R-HSA-209828, Tryptophan is hydroxylated | Category II | Serotonin |
|  | R-HSA-209859, Decarboxylation of 5-hydroxytryptophan forms serotonin | Category II | Serotonin |
|  | R-HSA-390666, Serotonin receptors | Category I | Serotonin |
|  | R-HSA-390648, Muscarinic acetylcholine receptors | Category I | Acetylcholine |
|  | R-HSA-209823, Tyrosine is hydroxylated to dopa | Category II | Dopamine |
|  | R-HSA-209924, Dopa is decarboxylated to dopamine | Category II | Dopamine |
|  | R-HSA-351596, Dopamine translocates to a secretory vesicle | Category II | Dopamine |
|  | R-HSA-390651, Dopamine receptors | Category I | Dopamine |
|  | R-HSA-209891, Dopamine is oxidised to noradrenaline | Category II | Norepinephrine |
|  | R-HSA-390696, Adrenoceptors | Category I | Norepinephrine |
|  | R-HSA-351604, Noradrenaline translocates to the cytosol | Category II | Epinephrine |
|  | R-HSA-209903, Noradrenaline is converted to adrenaline | Category II | Epinephrine |
|  | R-HSA-390696, Adrenoceptors | Category I | Epinephrine |

### 4) Gene list from the NeuronChat

Zhao et al. (2023) constructed a neural-specific curated list of cell-cell interactions. They curated a total of 373 entries consisted of eight neurotransmitters (dopamine, serotonin, glutamate, GABA, norepinephrine, epinephrine, acetylcholine, glycine), neuropeptide-mediated neurotransmission, synaptic adhesion, gap junctions, and gasotransmitters (NO and CO) into a database, namely NeuronChatDB. Their database consists of proteins that are related to the “ligand” of each cell-cell interaction and “receptor” proteins. We curated genes from their interaction databases by classifying genes that are related to the ligands, namely “ligand contributors”, as Category II, and genes that are related to the receptor functionalities as Category I in our work. The original NeuronChatDB is available from their original repository (https://github.com/Wei-BioMath/NeuronChatAnalysis2022/tree/main/NeuronChatDB_table).

**Supplementary information S2** LLM prompt for literature screening. [System prompt]

You are a biomedical literature reviewer. Classify whether the given ABSTRACT supports, refutes, or is neutral toward the HYPOTHESIS. Base your reasoning strictly on the ABSTRACT.

EVALUATION CATEGORIES:

- SUPPORT: The abstract provides evidence that the hypothesis is correct
- REFUTE: The abstract explicitly demonstrates the opposite direction
- NEUTRAL: The abstract shows no significant difference, unclear results, or does not address any phenomenon related to the hypothesis

CRITICAL CLARIFICATION:

- No significant difference or no difference shown → NEUTRAL (null finding, not refutation)
- Abstract doesn’t mention the phenomenon → NEUTRAL (no evidence to evaluate)
- Only classify as REJECT when an explicit opposite directional effect is shown

CONFIDENCE LEVELS:

- HIGH: Clear evidence directly addresses the hypothesis with explicit directional findings
- MEDIUM: Relevant findings present but with some ambiguity or indirect relevance
- LOW: Minimal evidence, borderline relevance, or high interpretive uncertainty

GUIDANCE FOR CONFIDENCE:

HIGH confidence applies when: The abstract explicitly reports directional effects with clear statistical significance or explicit null findings
MEDIUM confidence applies when: Findings of the abstract partially address the hypothesis with some ambiguity in interpretation
LOW confidence applies when: Abstract is tangentially related or confidence in the classification is uncertain

OUTPUT FORMAT (JSON):

~~~
{
  “verdict”: “SUPPORT” | “REFUTE” | “NEUTRAL”,
  “confidence”: “HIGH” | “MEDIUM” | “LOW”,
  “rationale”: “justification referencing the abstract, about 2-3 sentences“
}
~~~

[User prompt]

HYPOTHESIS: <HYPOTHESIS THAT USER WANTS TO EVALUATE>

TITLE: <PUBMED ARTICLE TITLE>

ABSTRACT: <PUBMED ARTICLE ABSTRACT>

Answer using JSON.

## Notes

### Competing Interest Statement

The authors have declared no competing interest.

